# AAV Kills Dividing Cells by Depleting PARP1 and Other DNA Damage Response Proteins

**DOI:** 10.1101/2025.10.14.682484

**Authors:** Sasha Friese, Junjie Zai, Grace Luzbetak, Nyonika Khanna, Johanna Gesperger, Chien-Hung Liu, Wynand P. Roos, Rasha Al-Rahahleh, Matisse Willardson, Feng Yang, Duoduo Fu, Yiming Han, Nadia Lintag, Jaiden Saykham, Joshua Le, Emeen Al-Delaimy, Nolan Soutipan, Ellen Duong, Jeremy Rich, Maria Carolina Marchetto, Michael Rosenfeld, Robert W. Sobol, Matthew Shtrahman

## Abstract

Recombinant adeno-associated virus (rAAV) is a replication-defective viral vector used in hundreds of human gene therapy trials, resulting in five FDA-approved therapies. Despite this success, rAAV-based gene therapies suffer from dose-limiting toxicities, resulting in several severe adverse reactions, including death. Previously, we discovered that rAAV rapidly kills mouse NPCs in vitro and in vivo. This vector contains a minimal genome comprised of 145-base pair inverted terminal repeats (ITRs) with a T-shaped hairpin structure that appears to be necessary and sufficient for this toxicity. However, the mechanism for AAV ITR toxicity is not known, and there have been few attempts to engineer ITRs to attenuate rAAV toxicity. In the current study, we explore the molecular mechanisms that drive dose-dependent rAAV toxicity in dividing human NPCs (hNPCs) and test whether disrupting these mechanisms mitigates this toxicity. Recombinant AAV infection induces aberrant cell cycle progression with activation of the ATM /CHK1/CHK2 pathway and expression of the DNA damage markers γH2AX and 53BP1. Affinity-based proteomics indicate that AAV ITRs bind to Poly-(ADP-Ribose)polymerase 1 (PARP1) and other DNA damage response (DDR) proteins involved in single-strand break repair (SSBR). Recombinant AAV infection attenuates poly-(ADP-ribose) (PAR) formation and mimics the antiproliferative effects of pharmacological PARP inhibitors used in cancer therapy. Moreover, treatment of hNPCs with PARP inhibitors is sufficient to reproduce many features of rAAV-induced toxicity. Finally, we demonstrate that eliminating the T-shaped hairpin within the AAV ITR reduces binding to SSBR proteins and the resulting rAAV toxicity. These findings suggest that rAAV infection induces replication stress and cell death in dividing hNPCs by functionally depleting PARP1 and other DDR proteins that are essential for DNA replication. This work fills substantial gaps in the understanding of the mechanisms of rAAV toxicity and has important implications for the development of safer rAAV-based human gene therapies.

**One Sentence Summary:** The rAAV genome binds to and depletes PARP1 and other SSBR proteins that are essential for DNA replication, resulting in DNA double stranded breaks, checkpoint activation, and cell death in dividing cells.

## Introduction

Recombinant adeno-associated virus (rAAV) is a replication-defective viral vector with no viral protein expression that is used widely in studies across mammalian biology and human gene therapy. Its broad tropism and stable transgene expression have made rAAV the vector of choice for hundreds of gene therapy trials, including five FDA-approved therapies. Despite this success, there are a growing number of experimental animal(*1–7*) and clinical studies(*8–11*) demonstrating severe adverse reactions caused by rAAV. Moreover, there have been at least 10 rAAV-related deaths reported since 2019(*12–17*). Unfortunately, the use of less common serotypes or engineered capsids to avoid severe immune reactions has not eliminated morbidity or mortality due to rAAV(*18*).

While rAAV-induced toxicity has been observed in a variety of cell types, dividing stem/progenitor cells appear to be particularly vulnerable. Previously, we discovered that rAAV ablates adult neurogenesis and induces apoptosis in dividing mouse neural progenitor cells (NPCs) *in vitro* and *in vivo* in a dose-dependent manner(*6*). In agreement with other studies^19,20^, this toxicity was attributed to the vector’s minimal genome, which contains a pair of palindromic 145 base-pair DNA inverted terminal repeats (ITRs) that are essential for multiple functions during the life cycle and the production of rAAV. These GC-rich DNA segments flank the transgene on both ends and contain a highly conserved T-shaped hairpin structure. Unfortunately, these essential roles and conserved features limit the ability to alter these sequences and fully characterize their contribution to rAAV toxicity. Despite these limitations, existing studies suggest that ITRs are both necessary and sufficient to explain many of the toxic effects observed after rAAV infection, which induces apoptosis and p53 activation in dividing cells(*6*, *19*). However, the upstream cellular pathways that are altered in this form of nucleic acid toxicity and why replicating cells are highly susceptible are not fully understood. Closing this knowledge gap is critical for minimizing dose-limiting toxicities of rAAV, particularly in children who are the most common recipients of rAAV-based gene therapies and have active proliferation of neural and other stem and progenitor cell populations.

The propensity of viruses to kill NPCs and other dividing cells is not unique to rAAV. Congenital infection by Zika virus, cytomegalovirus (CMV), Rubella, varicella zoster virus (VZV), herpes simplex virus (HSV), and human immunodeficiency virus (HIV) causes microcephaly through toxic effects on NPCs and impairment of neurogenesis(*21–25*). How these distantly related evolutionary viruses cause NPC toxicity at the molecular level and whether they might disrupt similar pathways is not known. In contrast, there is considerable evidence that loss-of-function mutations in DNA damage response (DDR) genes are a major driver of aberrant neurogenesis and cause heritable forms of microcephaly. This includes DDR proteins involved in DNA single-strand break repair (SSBR), which are critical for repairing DNA fragments during lagging strand synthesis and resolving stalled replication forks during S phase(*26–35*). However, aberrant DDR has not been a major focus of mechanistic studies investigating viral toxicity in NPCs and viral-induced microcephaly (exceptions include(*36*, *37*)), despite the established connection with heritable forms of the disease.

In the present study, we demonstrate that rAAV attenuates proliferation and kills dividing human embryonic stem cell-derived NPCs (hNPCs) and patient-derived glioblastoma stem cells (GSCs). The severity of this rAAV-induced toxicity correlates with the rate of cell proliferation. We also show that rAAV induces aberrant cell replication and activates CHK1, CHK2, and ATM, which are important regulators of cell cycle progression. In support of this, treatment with CHK2 inhibitors partially rescues rAAV-induced toxicity. In addition, replication stress induced by rAAV infection results in the formation of DNA breaks in the host genome as reported by increased numbers of γH2AX^+^ and 53BP1^+^ puncta, which do not colocalize with rAAV genomes. Our affinity-based proteomic experiments demonstrate that ITRs bind several DDR proteins, including Poly-(ADP-Ribose) polymerase 1 (PARP1) and other proteins involved in DNA single-strand break repair (SSBR). In addition, rAAV infection attenuates PAR formation and its toxic effects are additive with ABT-888, a pharmacological PARP inhibitor used in cancer therapy. Moreover, the effects of ABT-888 treatment in hNPCs mirror the molecular and cellular phenotypes of rAAV infection described above. Importantly, we show that eliminating the T-shaped hairpin of the ITR reduces binding to PARP1 and other SSBR proteins and mitigates rAAV toxicity in hNPCs. Collectively, these results indicate that during rAAV infection, AAV ITRs alter the cell cycle and induce cell death in hNPCs by binding to and functionally depleting DDR proteins that are essential for DNA replication. The current study establishes the mechanistic framework for developing safer rAAV-based human gene therapies and the foundation for harnessing rAAV toxicity to engineer new oncolytic viruses in the future.

## Results

### rAAV induces potent dose-dependent toxicity in hNPCs, which correlates with the rate of cell proliferation

Wild-type AAV is a human pathogen. However, most rAAV serotypes exhibit broad tropism across mammalian cells. Consistent with this, we had previously shown that rAAVs, containing AAV2 ITRs, attenuate proliferation and kill dividing mouse NPCs both *in vitro* and *in vivo*, independent of AAV capsid serotype(*6*). To explore mechanisms of rAAV toxicity in an NPC model more relevant for human gene therapy and microcephaly, we tested the effect of rAAV infection on embryonic stem-cell-derived hNPCs. For these experiments, we utilized an rAAV1 (with AAV2 ITR) encoding a Cre-dependent GFP in non-Cre-expressing cells to minimize contributions of protein expression to vector toxicity. Human NPCs were labeled with the fluorescent cell viability marker propidium iodide (PI) and the confluence (phase contrast) and cell death (fluorescence) were imaged every 4 hours for approximately 72 hours. An example experiment is described in Figure 1, where hNPCs are plated into a 96-well plate and infected with rAAV at three different multiplicities of infection (MOIs): 1E4, 1E5, and 1E6 viral genome copies (gc)/cell and saline control. Recombinant AAV infection resulted in a robust dose-dependent reduction in cell proliferation and cell death (Figs. 1A and 1B, respectively). A similar level of toxicity was observed when rAAV infected patient-derived glioblastoma stem cells (Fig. S1). Figures 1D and 1E quantify rAAV dose-dependent reduction in proliferation and increase in cell death across 16 different experiments, each performed in triplicate. The average reduction in proliferation at MOI = 1E4, 1E5, and 1E6 genome copies (gc)/cell was 22 ± 2%, 63 ± 2%, and 76 ± 2%, respectively; the average increase in cell death at these MOIs was 80 ± 5%, 350 ± 20%, and 490 ± 20%, respectively. For a given MOI, hNPCs exhibit significant heterogeneity in their sensitivity to rAAV infection (total coefficient of variation (cv) for reduction in proliferation = 65.1%, 23.4%, 14.2% and cv for increase in cell death = 43.3%, 40.8%, 34.0% calculated over all wells at MOI = 1E4, 1E5, and 1E6 gc/cell, respectively). By comparison, rAAV toxicity measured in wells from the same experiment (identical vial and passage) generally exhibited less variance in sensitivity, particularly at high MOI (Fig. S2, average within-experiment cv for reduction in proliferation = 124.0%, 3.5%, 1.2% and average cv for increase in cell death = 16.3%, 6.5%, 5.2% calculated over all wells within the same experiments as above). Another indication that this heterogeneity in rAAV toxicity is not primarily due to random experimental variability is that the reduction in proliferation and the increase in cell death for cells in the same well were tightly correlated (Fig. 1F, Spearman coefficient r = 0.96, p < 0.0001).

**Figure 1.**
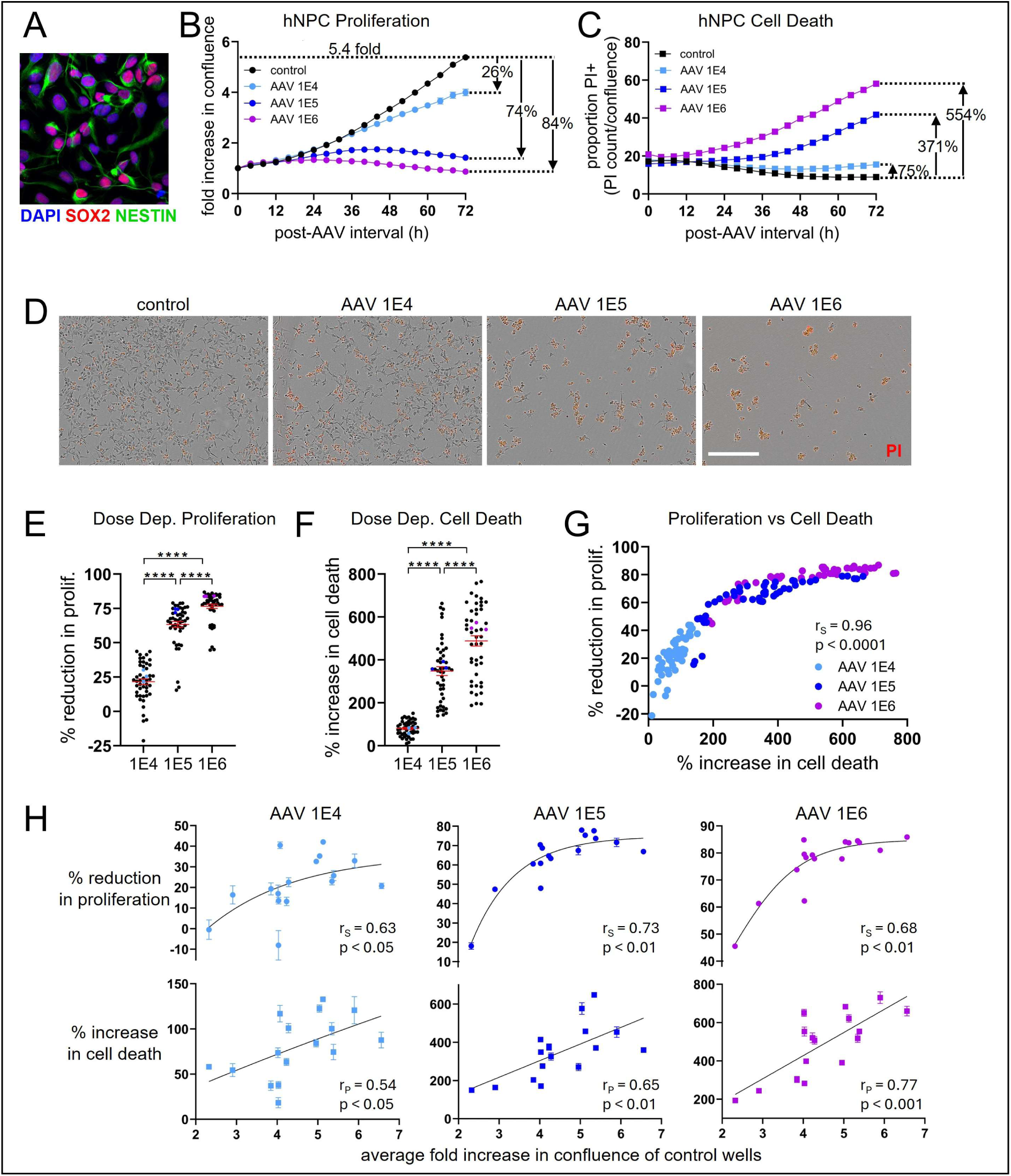
Recombinant AAV induces strong dose-dependent toxicity in hNPCs that increases with the rate of proliferation. **(A)** Example immunocytochemical staining of hNPCs expressing SOX2 and NESTIN (**B**) Example cell confluence measurements from time-lapse microscopy of hNPCs plated on a 96-well plate and infected with rAAV at various MOIs and saline control demonstrate strong dose-dependent attenuation of cell proliferation. Each point represents normalized confluence averaged over n = 3 wells. (**C**) Dose-dependent rAAV-induced cell death is measured in the same wells by calculating the proportion of PI+ hNPCs. (**D**) Representative images showing confluence (brightfield) and PI labeling (red) of hNPCs 72 hours post-viral transduction at MOIs of 10^4^, 10^5^, and 10^6^, and control. Scale bar = 400 µm (**E**) Reduction in cell proliferation and (**F**) increase in cell death at 72 hours for each well (n = 48) are plotted for each MOI and show dose dependence and significant variability between experiments (one-way ANOVA followed by pairwise comparison using Tukey’s test). Wells from the experiment described in (B) and (C) are marked in color. (**G**) Increase in cell death is plotted versus reduction in proliferation for each well, demonstrating a strong correlation between these metrics. (**H**) Average reduction in proliferation (*top*) and increase in cell death (*bottom*) for each experiment (n = 3 wells per experiment) are plotted versus the average increase in confluence of the control wells for the same experiment demonstrating that the magnitude of rAAV toxicity increases with the rate of proliferation at all MOIs. The black line represents the best fit to the exponential plateau function y = a + b*exp(-k*x) or linear regression for reduction in proliferation and increase in cell death, respectively. All error bars represent standard error of the mean. Error bars smaller than points are not shown. Nonlinear Spearman (r_S_) and linear Pearson (r_P_) correlation coefficients; p < 0.05, 0.01, 0.001, 0.0001 depicted respectively by *, **, ***, ****.

Previously, we had shown that adult mouse NPCs that label with BrdU or express the transient amplifying NPC marker Tbr2 are significantly more susceptible to rAAV toxicity compared to other adult NPC populations *in vivo*, suggesting that rAAV preferentially kills dividing cells(*6*). We hypothesized that the differences in cell proliferation rates between experiments, which reflect the fraction of replicating cells or mitotic index, could explain the observed heterogeneity in rAAV toxicity. To quantify the rate of proliferation, we calculated the average fold increase in confluence at 72 hours for the control wells receiving saline for each experiment. We plotted this metric of proliferation against rAAV-induced attenuation in proliferation and increase in cell death in the same experiment. Figure 1G shows that the susceptibility to rAAV toxicity increases monotonically with increasing rate of hNPC proliferation, with higher MOIs showing less variability (Spearman coefficient r_S_ = 0.63, 0.73, 0.68 and p-value < 0.05, 0.01, 0.01 for reduction in proliferation and Pearson coefficient r_P_ = 0.54, 0.65, 0.77 and p-value < 0.05, 0.01, 0.001 for increase in cell death calculated for MOI = 1E4, 1E5, and 1E6 gc/cell, respectively). This correlation between the magnitude of rAAV toxicity and the rate of proliferation is consistent with the hypothesis that rAAV infection disrupts cell division.

### rAAV infection induces aberrant cell cycle progression and activation of CHK1, CHK2, and ATM

Based on the above findings, we hypothesized that rAAV infection is a source of replication stress, inducing aberrant cell cycle progression and subsequent cell death in dividing cells. To test this assumption, we infected hNPCs with rAAV at an MOI of 1E6 and stained the cells with DAPI 24 hours later, an early time point at which growth curves begin to demonstrate a clear decrease in proliferation. At 24 hours post-infection, rAAV-infected hNPCs demonstrate nuclear enlargement (control 111 ± 2 µm^2^ vs rAAV 152 ± 3 µm^2^, p < 0.0001), consistent with aberrant cell cycle progression (Figs. 2A and 2B)(*38–41*). Furthermore, nuclear enlargement displayed dose dependence when hNPCs were infected with rAAV at different MOIs within a single experiment (Fig. 2C). However, cell cycle analysis in hNPCs shows that neither infection with rAAV, nor treatment with the antimitotic agent Nocodazole, induce substantial cell cycle arrest, indicating that apoptosis may be more prevalent at these doses (Figs. 1, S3, and S4).

**Figure 2.**
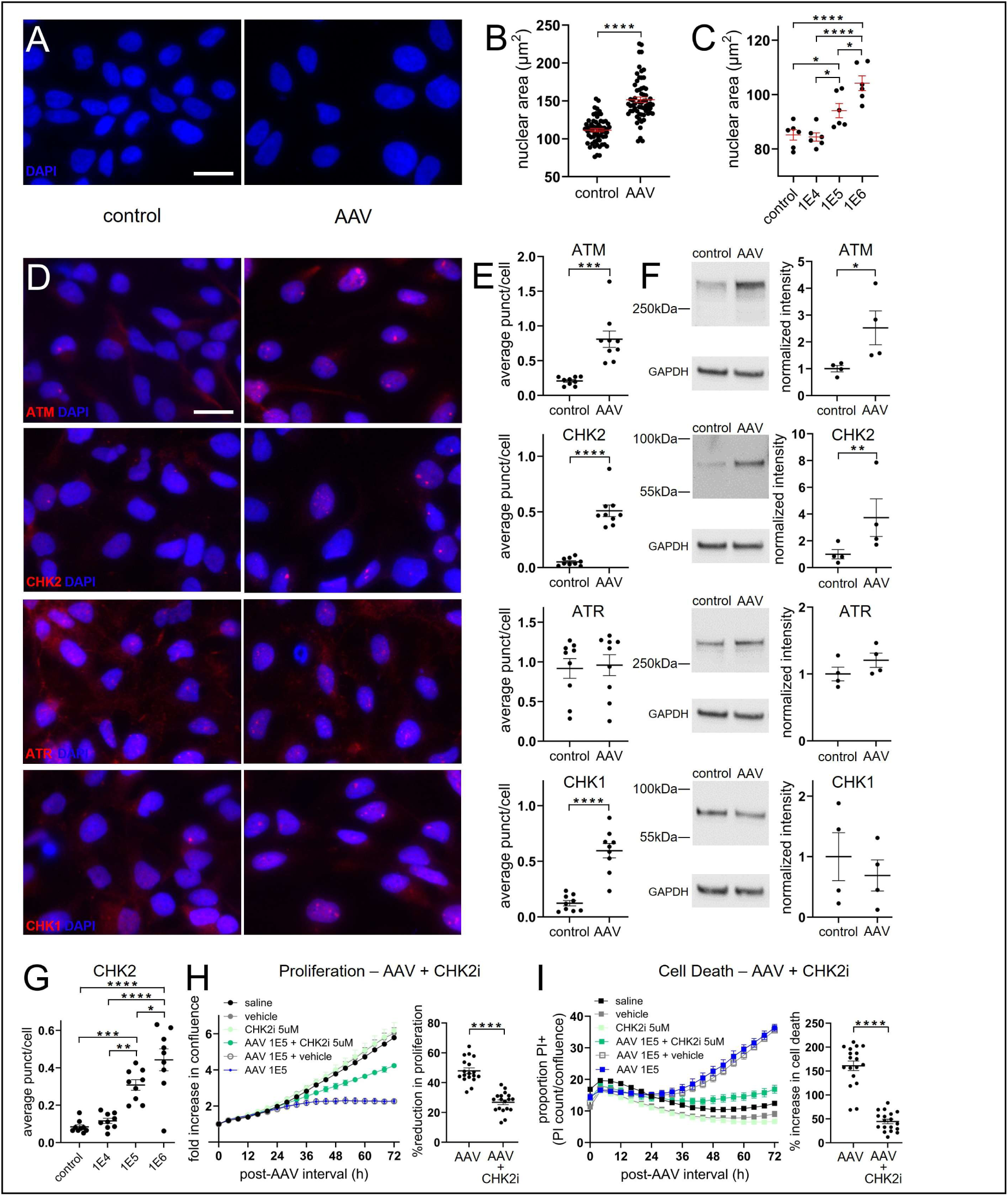
rAAV infection induces aberrant cell cycle progression and activation of the ATM/Chk1/Chk2 pathway. (**A**) DAPI fluorescent image showing an extreme example of nuclear enlargement 24 hours after treatment with rAAV (MOI = 1E6, *right*) compared to control (*left*). **B**) Average nuclear area of cells treated with rAAV compared to control cells (n = 63 wells per condition; unpaired t-test). (**C**) Cells treated with different doses of rAAV within a single experiment demonstrate dose-dependent nuclear enlargement (n = 6 wells per condition; one-way ANOVA followed by pairwise comparison using Tukey’s test). (**D**) Representative ICC images of control (*left*) and rAAV-infected (*right*) hNPCs stained for ATM, CHK2, ATR, and CHK1 24 hours after infection. Scale bar = 20 µm. (**E**) The average number of ATM+, CHK2+, ATR+, and CHK1+ foci per nucleus are compared for wells infected with rAAV versus control (n = 9 wells per condition; unpaired t-test). (**F**) Total protein expression in hNPCs infected with rAAV was measured by western analysis and compared to control conditions (n = 4 experiments; ratio paired t-test). All values are normalized by the average control protein expression. (**G**) The average number of CHK2+ foci per cell is plotted at each MOI, exhibiting a dose-dependent response (n = 9 wells per condition; one-way ANOVA followed by pairwise comparison using Tukey’s test). (**H**) *Left* Example of partial rescue of rAAV-induced reduction in proliferation in hNPCs treated with and without the CHK2 inhibitor (n = 3 wells per condition). *Right*: Reduction in hNPC proliferation at 72 hours is plotted for wells infected with rAAV and compared to those treated with both rAAV and CHK2 inhibitor (n = 18 wells per condition; unpaired t-test). (**I**) *Left*: Example of partial rescue of rAAV-induced cell death in hNPCs treated with and without the CHK2 inhibitor (n = 3 wells per condition). *Right*: Quantification of cell death at 72 hours is plotted for wells infected with rAAV and compared to those treated with both rAAV and CHK2 inhibitor (n = 18 wells per condition; unpaired two-tailed t-test).

**Figure 3.**
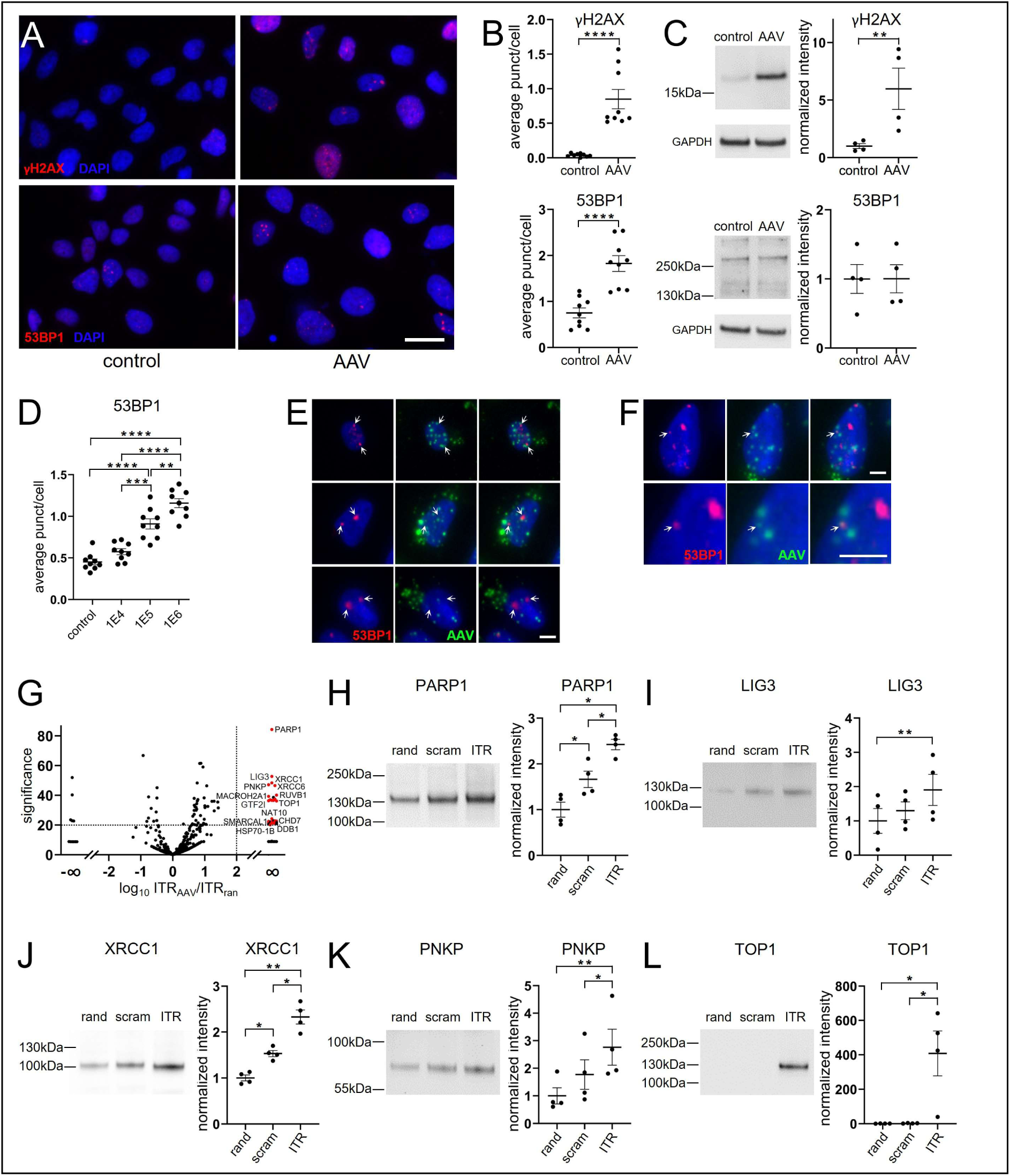
rAAV induces double-stranded DNA breaks and recruits host DDR proteins. (**A**) Representative ICC images of control (*left*) and rAAV-infected (*right*) hNPCs stained for γH2AX and 53BP1 24 hours after treatment. Scale bar 20 µm. (**B**) The average number of γH2AX+ and 53BP1+ foci per nucleus is compared for wells infected with rAAV for 24 hours versus control (n = 9 wells per stain at each condition; unpaired t-test). (**C**) Total protein expression of γH2AX+ and 53BP1+ in cells infected with rAAV are measured by western analysis and compared to control conditions (n=4; ratio paired t-test). All values are normalized by the average control protein expression. (**D**) The average number of 53BP1+ foci per cell is plotted at each MOI and exhibits a dose-dependent response (n = 9 wells per condition; one-way ANOVA followed by Tukey test). (**E**) Representative images of green foci marking rAAV genomes via in situ hybridization with RNAscope probes for the EGFP gene and red 53BP1+ foci exhibit no overlap within hNPCs. Scale bar 5 µm. (**F**) Low (*top*) and high (*bottom*) magnification images of a rare example of overlap between a single rAAV genome+ focus (green) and 53BP1+ focus (red). (**G**) Relative abundance of nuclear proteins isolated by AAV2 ITR (“ITR_AAV_”) compared to a random 145-base pair single-stranded DNA sequence (“ITR_RAN_”) via an affinity pull-down assay in glioblastoma stem cells and measured by mass spec. The top 32 proteins (defined as significance ≥ 20, log_10_(ITR_AAV_/ITR_RAN_) ratio ≥ 2) pulled down by ITR_AAV_ were not detectable in the ITR_RAN_ fraction, yielding a log_10_(ITR_AAV_/ITR_RAN_) ratio = ∞. Twenty-five of the top 32 proteins participate in DDR (marked with red points) and 14 of these DDR proteins interact with PARP1 and are labeled. (**H-L**) Repeat of affinity pull-down assay in hNPCs followed by western analysis confirms that PARP1 and PARP1-associated proteins identified in (H) are preferentially bound by AAV2 ITR compared to scrambled and random control sequences (repeated measures one-way ANOVA of log_10_[protein expression] followed by Tukey test).

To explore the mechanism by which rAAV alters the cell cycle, we tested to what extent the ATM/CHK2 and the ATR/CHK1 pathways are activated in hNPCs in response to rAAV infection. Although these critical pathways interact, they can each be preferentially activated by different stressors during cell division(*42*). ATM and CHK2 showed consistent activation in response to rAAV infection, evident from both an increased number of phosphorylated ATM^+^ and CHK2^+^ puncta (Fig. 2E) and increased total expression of these proteins by western blot analysis (Fig. 2F). In addition, the degree of checkpoint activation, measured via immunocytochemistry by the number of CHK2^+^ puncta, displayed a clear dose dependence and was proportional to MOI (Fig. 2G). In contrast, there was no evidence of ATR activation in response to rAAV, while CHK1 activation was evident only from the increased number of CHK1^+^ puncta (Fig. 2E,F). Finally, when low-dose treatment with the CHK2 inhibitor (CHK2i) BML-277 was combined with rAAV infection, it partially rescued rAAV-induced toxicity in hNPCs (reduction in proliferation: rAAV 48 ± 2% vs rAAV + CHK2i 27 ± 2%, p < 0.0001; increase in cell death: rAAV 161 ± 10% vs rAAV + CHK2i 45 ± 5%, p < 0.0001, Figs. 2H and 2I). Collectively, these results indicate that rAAV infection is toxic to replicating cells, inducing replication stress and altering cell cycle progression in a dose-dependent manner via aberrant checkpoint activation, particularly via the ATM/CHK2 pathway.

### rAAV induces double-stranded DNA breaks and recruits host DDR proteins

The canonical trigger for activation of ATM/CHK is DNA damage in the form of double-stranded breaks. Infection with wild-type AAV, which contains the full viral genome and integrates into the host genome, has been shown to induce the DDR in the host(*43–45*). However, the extent to which rAAV, which alone has limited capacity to integrate, causes a similar DDR response, is not fully established(*46–48*). To explore whether rAAV-induced replication stress can induce DDR, we infected hNPCs with rAAV (1E6 gc/cell) and stained for the DDR marker γH2AX 24 hours after infection (Fig. 3A). Infected hNPCs exhibited a statistically significant increase in the number of γH2AX^+^ nuclear puncta (Fig. 3B) and total expression as measured by western blot analysis (Fig. 3C). Not all hNPC nuclei with increased γH2AX expression exhibited clear γH2AX^+^ puncta formation, possibly due to the difficulty in resolving individual puncta in the setting of dense γH2AX activation. Alternatively, increased γH2AX expression may be diffuse and not always indicative of DNA breaks(*49*). To address this, we stained hNPCs for 53BP1, a marker more specific for double-stranded DNA breaks(*50*). Twenty-four hours after infection, cells exhibited a marked increase in the number of 53BP1^+^ foci (Figs. 3A and 3B), with no change in total expression (Fig. 3C). Importantly, the overwhelming majority (98.8 ±0.7%; n=6 wells) of 53BP1^+^ puncta (total n=417) did not colocalize with the rAAV genome as measured by a combination of immunocytochemistry and *in situ* hybridization with an RNAscope probe that hybridizes to the rAAV GFP transgene. The above is consistent with the hypothesis that the DNA breaks recruiting 53BP1 are largely chromosomal and located within the host genome (Figs. 3E and 3F).

DDR is initiated by activation of upstream DDR sensors that identify DNA breaks, stalled replication forks, hairpins, G-quadruplexes, or other noncanonical DNA structures. We previously demonstrated that the impact of rAAV on cell proliferation and survival is rapid and can be observed as early as 12 hours after infection *in vivo*(*6*). This led us to hypothesize that the upstream sensor of AAV ITRs is normally present and surveys the nuclear genome, and therefore does not require *de novo* transcription or translation in response to rAAV infection. To identify candidate ITR-binding proteins, we performed an affinity pull-down assay to isolate nuclear proteins that preferentially bind synthetic biotinylated AAV2 ITR compared to a random 145-base-pair single-stranded DNA sequence. We used the patient-derived glioblastoma stem cells described above, which can be propagated easily and are sensitive to rAAV-induced toxicity (Fig. S1). PARP1, a first responder in the DDR, was the candidate protein with the largest significance and relative abundance (ITR_AAV_/ITR_ran_ ratio). PARP1 yielded 33 unique peptides in the AAV2 ITR isolate, whereas no PARP1 peptides exceeding the quality threshold were detected in the random-ITR fraction (Fig. 3H). The 4 most significant and abundant candidate ITR-binding proteins PARP1, LIG3, XRCC1, and PNKP are core proteins involved in DNA SSBR(*51–53*) (Fig. 3G). In addition, the candidate protein TOP1 is critical for unwinding DNA during replication and is known to interact with PARP1 during replication fork stalling(*54*, *55*). Notably, loss-of-function mutations in genes that code for these proteins have been associated with microcephaly and early neurodegeneration(*56–60*). Moreover, 24 out of the top 32 candidate AAV2 ITR-binding proteins, defined as having log ratio ≥ 2 and significance ≥ 20, have been implicated in the DDR (Supplementary Table 1). Fourteen of these proteins are known to interact with PARP1. We then repeated these experiments in primary mouse NPCs from E15.5 mice. Ten of the top candidate AAV2 ITR-binding proteins were identified in both experiments, including all the SSBR core proteins listed above and TOP1 (Fig. S5, Supplementary Table 2).

We went on to test the affinity of AAV2 ITR for PARP1, LIG3, XRCC1, PNKP, and TOP1 in hNPCs, using a similar affinity pull-down assay that also included a scrambled ITR sequence. The scrambled ITR sequence controls for the high GC content of the AAV2 ITR, but contains no T-shaped hairpins as predicted by the mFold algorithm(*61*) (Fig. S6). These experiments demonstrated that PARP1 and PARP1-associated proteins bind to AAV2 ITR > scrambled ITR > random sequence. Taken together, these data support the notion that rAAV recruits host DDR machinery important for DNA replication, which causes replication stress and subsequent DNA double-strand breaks in the host genome.

### rAAV toxicity is not rescued by NAD^+^ supplementation

PARP1 is an upstream sensor for damaged DNA and noncanonical DNA structures, such as hairpins and stalled replication forks(*62*, *63*). These DNA structures trigger activation of PARP1, which catalyzes the formation of poly-(ADP-ribose) (PAR). PARP1 catalyzes covalent binding of this polymer to itself, SSBR core proteins, and other DDR proteins in a posttranslational modification known as PARylation. Overactivation of PARP1 depletes NAD^+^, the major substrate for PAR formation, leading to ATP depletion, energetic collapse(*64–66*), and impairment of critical cellular functions such as glycolysis(*67*) and cell division(*26*). Supplementation with NAD^+^ precursors, such as nicotinamide riboside (NR), have been shown to rescue overactivation of PARP1 in a number of experimental contexts(*66*, *68*, *69*). One plausible mechanism driving rAAV toxicity is that either ITR hairpins or chromosomal DNA breaks consequent to rAAV infection over activate PARP1, leading to NAD^+^ depletion and energetic collapse in hNPCs. To probe this hypothesis, we supplemented infected hNPCs cells with NR to test if preservation of NAD^+^ levels can attenuate rAAV toxicity (Fig. S7A and S7B). Time-lapse microscopy measurements of cell confluence and viability with and without NR supplementation show no rescue of rAAV toxicity at all MOIs (reduction in proliferation: rAAV 16 ± 2%, 58± 4%, 79± 3% vs rAAV + NR 20 ± 2%, 63± 3%, 79± 3%, n.s.; increase in cell death: rAAV 60 ± 10%, 230± 30%, 460± 60% vs rAAV + NR 59 ± 10%, 260± 30%, 450± 60%, n.s. at MOIs 1E4, 1E5, and 1E6, respectively), suggesting that rAAV-induced toxicity is not mediated by PARP1 overactivation.

### rAAV-induced toxicity mimics pharmacological inhibition of PARP1

PARP1, together with other SSBR core proteins, serve multiple functions during replication, including lagging strand synthesis and resolution of stalled replication forks, and is important for proper progression through the cell cycle(*26–35*). In addition, PARP1 activity rises with increasing cell proliferation (*70*, *71*). Pharmacological PARP inhibitors, which trap PARP1 on the DNA, attenuate proliferation and kill cancer cells in a variety of experimental and clinical contexts(*72*). We hypothesized that binding of AAV ITRs to PARP1 and other SSBR core proteins would mimic pharmacological inhibition of these proteins. To explore whether rAAV inhibits PARP1 activity, we employed a cellular assay that utilizes an engineered EGFP-fusion PAR binding protein (LivePAR) to quantify the formation of PAR in response to genotoxic agents in U2OS cells(*73–75*). To induce substantial and stable PAR formation, we utilized a combination of the genotoxic agent MNNG, the NAD^+^ supplement NRH, and the PAR glycohydrolase inhibitor (PARGi) PDD00017273, which prevents break down of PAR (Figs. 4A and 4B)(*76*). Consistent with previous studies(*73–75*), we observed robust formation of nuclear EGFP-tagged PAR^+^ puncta induced by these agents. This effect was nearly abolished by pre-treatment with ABT-888, a potent inhibitor of PARP1 and PARP2. Infecting cells with rAAV 24 hours before MNNG/NRH/PARGi treatment also attenuated PAR formation, although to a lesser extent than the PARP inhibitor. Our findings support the hypothesis that AAV ITRs not only bind to PARP1, but also inhibit its activity.

**Figure 4.**
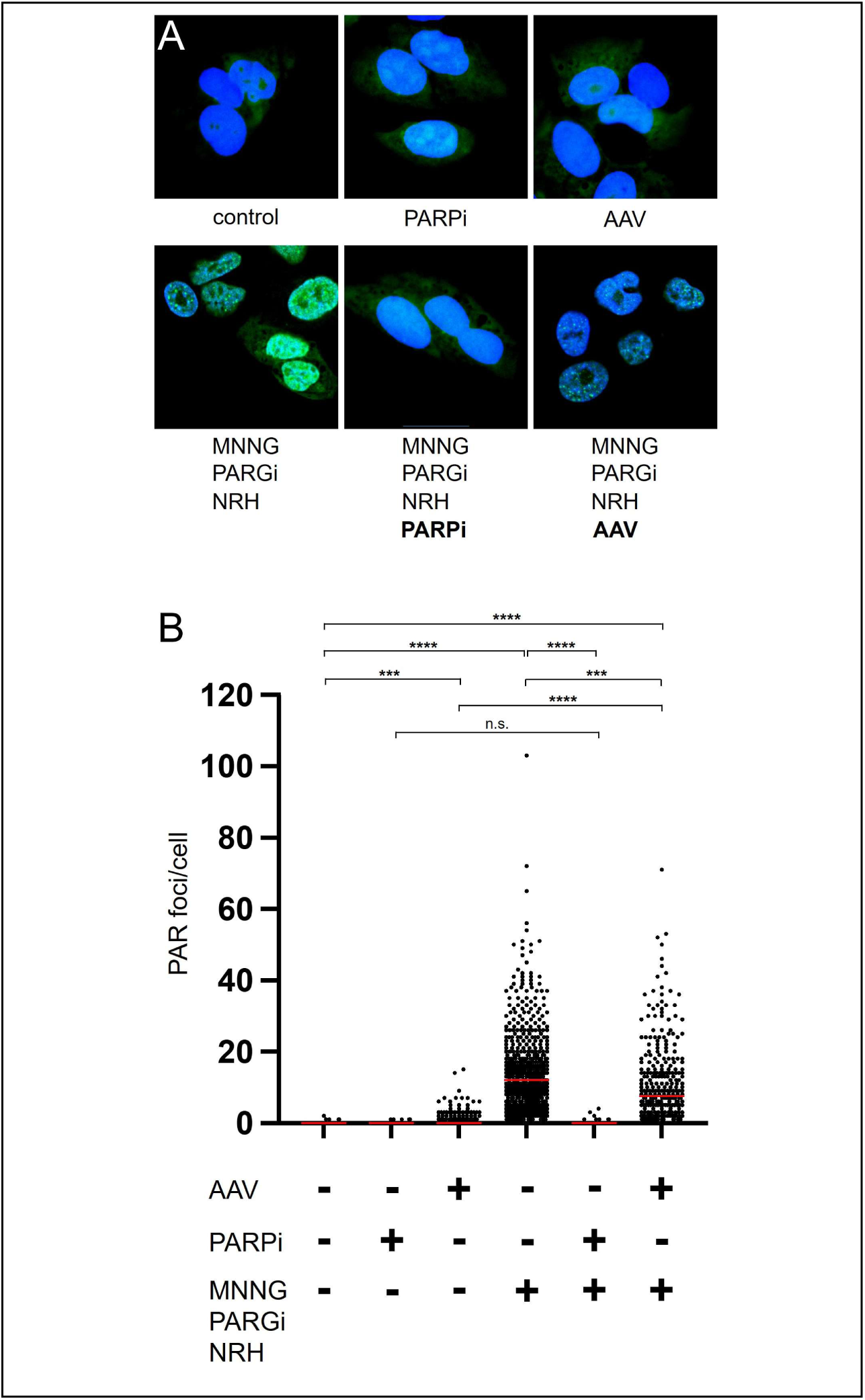
rAAV attenuates PAR formation in response to genotoxic agents. (**A**) *Left*: Representative fluorescent confocal microscopy images of US02 cells expressing LivePAR, an engineered EGFP-fusion PAR-binding protein that localizes to PAR+ puncta in response to treatment with MNNG/NRH/PARGi (*bottom*), which induces substantial genomic stress compared to control conditions (*top*). *Middle*: Formation of EGFP+ puncta by MNNG/NRH/PARGi (*bottom*) are abolished by pretreatment with the PARP inhibitor ABT-888 (*top*). *Right*: Pretreatment with rAAV (*bottom*) attenuates, but does not completely prevent, formation of EGFP+ puncta formation in response to MNNG/NRH/PARGi treatment compared to control conditions (*top*). (**B**) Quantification of the number of GFP+ puncta under different treatment conditions displayed below the graph demonstrates the complete and partial attenuation of MNNG/NRH/PARGi-induced EGFP+ puncta by ABT-888 and rAAV, respectively; (Kruskal-Wallis followed by Dunn’s multiplecomparisons test).

To further test this hypothesis, we asked whether treatment of hNPCs with a PARP inhibitor results in a similar dose-dependent toxicity as rAAV infection. HNPCs were treated with ABT-888 and imaged as described in Figure 1. Figures 5A-D show that ABT-888, but not vehicle alone, attenuated proliferation and killed hNPCs in a dose-dependent manner like rAAV infection. We surmised that if both rAAV and ABT-888 exhibit dose-dependent effects and work through similar mechanisms, simultaneous treatment with modest doses of both should yield an additive effect, while the combined toxicity at higher doses should saturate, similar to treatment with each agent individually. To test this assumption, we added varying doses of ABT-888 to rAAV at an MOI of 1E4, which alone has modest and variable toxicity (Figs. 1D and 1E), and measured reduction in hNPCs proliferation and viability. An example of the additive effects of these agents is shown in Figures 5D and 5E. As expected, 1E4 rAAV enhanced ABT-888-induced cell toxicity, with a linear additive effect at low doses and saturation at higher doses of ABT-888 (Figs. 5G).

**Figure 5.**
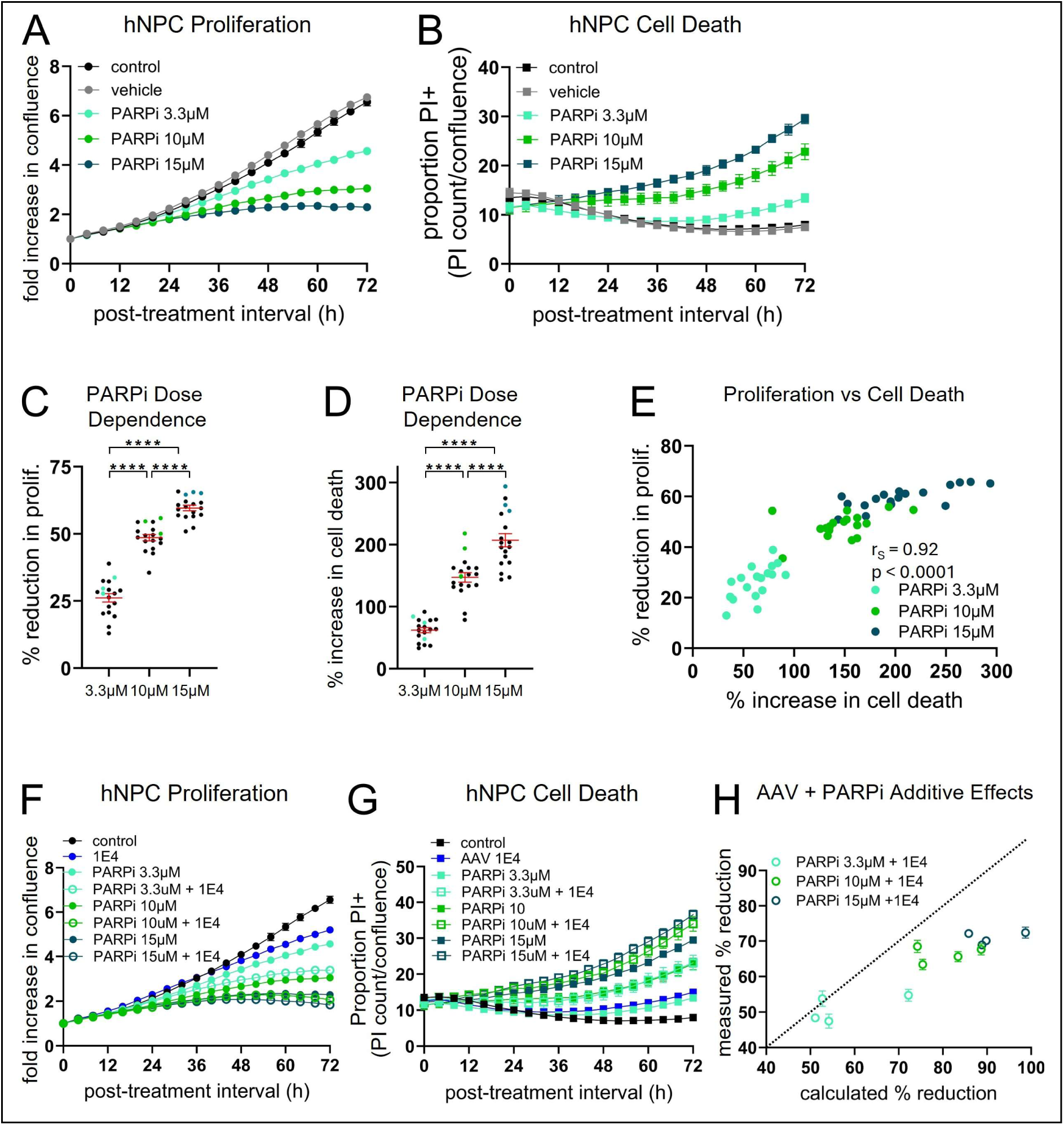
PARP inhibitor treatment induces dose dependent toxicity in hNPCs that is similar to and additive with rAAV infection. (**A**) Example measurements from time lapse microscopy treated with ABT-888 at various doses in addition to saline and vehicle control demonstrating strong dose dependent attenuation of cell proliferation and (**B**) cell death in hNPCs (n = 3 wells per condition) (**C**) Reduction in cell proliferation and (**D**) increase in cell death at 72 hours for each well (n = 18) is plotted for each dose and shows strong dose dependence (one-way ANOVA followed by pairwise comparison using Tukey’s test). Wells from the example experiment displayed in (A) and (B) are marked in color. (**E**) Reduction in cell proliferation is plotted versus increase in cell death for each well, demonstrating a strong correlation between these metrics using nonlinear Spearman (r_s_) coefficient. (**F**) The same experiment described in (A) and (B) is shown quantifying the effects of PARP inhibitor with (open circles) and without (closed circles of the same color) the addition of rAAV 1E4 on hNPC proliferation and (**G**) cell death. Data summarizing the additive effects of these agents is plotted in (**H**), where y-axis plots the reduction of proliferation measured when combining these agents experimentally and the x-axis plots the linear sum of the reduction in proliferation calculated from wells receiving each of these agents separately. Dotted line represents y = x.

We next asked if ABT-888 treatment induces nuclear enlargement and activates checkpoint pathways analogous to the action of rAAV discovered in our experiments. Human NPCs treated with ABT-888 for 24 hours exhibited enlarged nuclei, but not substantial cell cycle arrest, similar to rAAV infection (Figs. 6A, 6B, and S4). Treating hNPCs with ABT-888 resulted in consistent activation of ATM^+^, CHK2^+^, and CHK1^+^, but not ATR^+^, puncta, nearly identical to the pattern observed with rAAV infection (Figs. 6C and 6D). We also tested whether treatment with ABT-888 for 24 hours induces DNA damage in hNPCs. Again, similar to rAAV infection, robust expression of γH2AX and 53BP1^+^ puncta were observed in hNPCs treated with ABT-888 (Figs. 6C and 6D). Collectively, these data suggest that toxicity induced by rAAV infection mimics the action of PARP inhibition in hNPCs.

**Figure 6.**
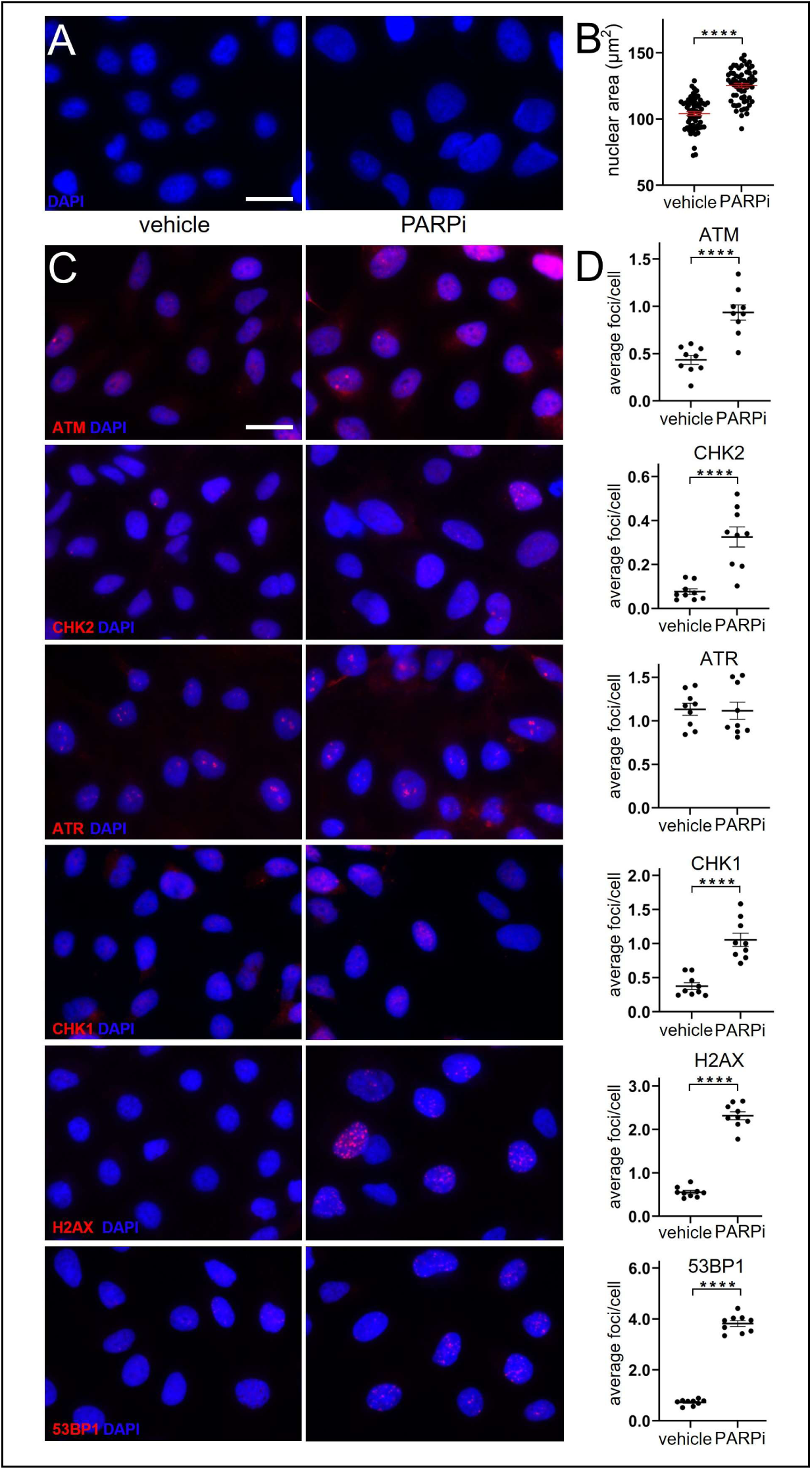
PARP inhibitor treatment induces aberrant cell cycle progression and activation of the ATM/pChk2 pathway. (**A**) DAPI fluorescent image showing an extreme example of nuclear enlargement 24 hours after treatment with 15 μM of the PARP inhibitor ABT-888 (*right*) compared to the control (*left*). **B**) Average nuclear area (n = 63 wells per condition; unpaired t-test) of cells treated with 15 μM ABT-888 compared to control cells. (**C**) Representative ICC images of control (*left*) and PARP inhibitor-treated (*right*) hNPCs stained for ATM, CHK2, ATR, CHK1, γH2AX, and 53BP1 24 hours after treatment. Scale bar 20 µm. (**D**) The average number of ATM+, CHK2+, ATR+, CHK1+, γH2AX+, and 53BP1+ puncta per nucleus is calculated (using the same parameters as used for rAAV described in Figure 2) and compared for wells infected with PARP inhibitor for 24 hours versus control (n = 9 wells per condition; unpaired t-test).

### AAV ITR T-shaped hairpin binds PARP1 and PARP1-associated proteins and is necessary for rAAV toxicity

DNA with non-canonical secondary structure, such as DNA hairpins, encountered during DNA replication can impede or even stall replication fork progression(*77–79*). PARP1, TOP1, and other DDR proteins are responsible for recognizing these structures and resolving stalled replication forks. PARP inhibitors are thought to kill dividing cells by binding and trapping PARP1 bound to DNA, functionally depleting this important protein and inducing insurmountable replication stress and subsequent cell death(*80*). We hypothesized that the T-shaped hairpin secondary structure of the AAV ITR binds and sequesters PARP1 and other DDR proteins and induces toxicity in dividing cells in a manner analogous to PARP inhibitors. To test this hypothesis, we synthesized an rAAV9 that expresses EGFP and contains mutant ITRs, where the B/B’ and C/C’ hairpins have been deleted, leaving only a minimal hairpin(*81*) (ΔBC ITR, Fig. 7A). Recombinant AAV with the ΔBC ITR mutation has been shown to produce 8-fold lower viral titers compared to viruses with AAV2 ITR. Given the steep dose-dependent toxicity of rAAV described above (see Fig. 1), accurate titers are essential for characterizing viral toxicity. Thus, we utilized three different independent methods to compare titers between the mutant and wt rAAVs. First, qPCR measurements confirmed the 8-fold reduction in the number of ΔBC viral genomes (5 E12 gc/ml) relative to an rAAV9 control containing an identical construct, but with wild-type (wt) AAV2 ITRs (4 E13 gc/ml; Table 1). We then performed transmission electron microscopy to compare the number of full and empty capsids in wt and ΔBC mutant rAAV9 (Fig. 7B, Table 1). A third rAAV9 carrying the same EGFP transgene (Addgene) with a known titer obtained via digital droplet PCR served as an additional control for relative and absolute titers (Table 1). The number of full capsids in these three viral preparations again confirmed the 8-fold reduction in the ΔBC rAAV9 titer compared to the wt control rAAV9 titer. In addition to the relative reduction, the experiment also confirmed the absolute titers obtained by qPCR, which consistently differed from digital droplet PCR by a factor of 2, as previously described (Table 1)(*82*). Lastly, we measured the total number of capsids in these three rAAV preparations using ELISA against the capsid protein, which agreed with the total (full + empty) capsid titers measured by electron microscopy (Table 1).

**Figure 7.**
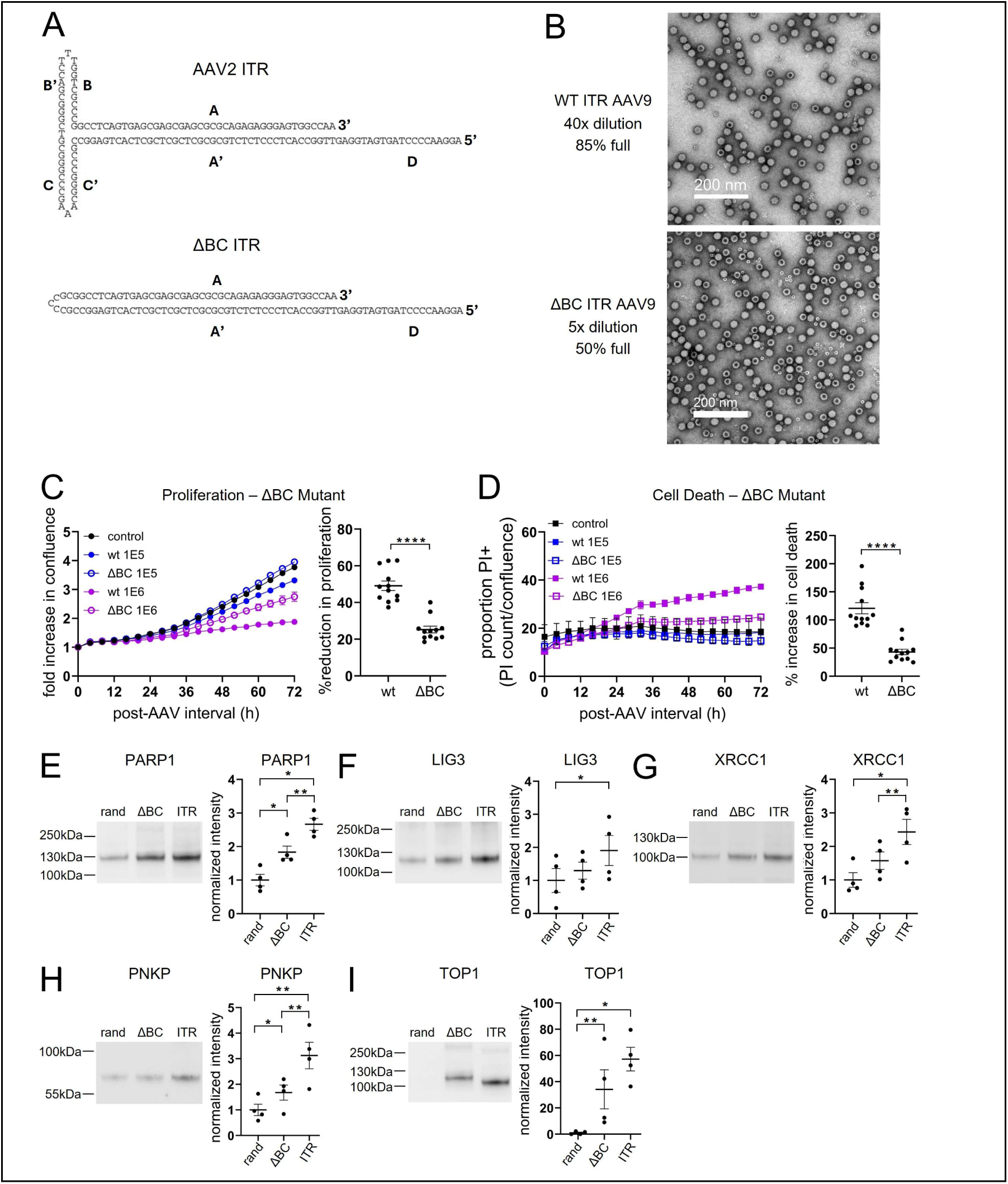
AAV ITR T-shaped hairpin binds PARP1 and PARP1-associated proteins and is necessary for rAAV toxicity. (**A**) Schematic of the wildtype (wt) AAV2 ITR (*top*) and the ΔBC ITR mutant (*bottom*) that lacks the B/B’ and C/C’ hairpins. (**B**) Representative TEM images of wt and mutant rAAVs showing full and empty capsids. Empty capsids collapse and contain pooled contrast giving a donut- or ring-like appearance. TEM allows for quantification of the relative viral titer, which can be compared not only between the wt control and ΔBC mutant, but also to an rAAV with known titer measured by another method (see Results, Table 1). Scale bar 200 nm. (**B**) Example and summary data for (**C**) cell proliferation and (**D**) cell death measurements from time-lapse microscopy of hNPCs infected with rAAVs encoding the ΔBC ITR demonstrate decrease toxicity compared to wt rAAV control at the same MOI (n = 12 wells per condition; unpaired t-test). (**E-I**) Affinity pull-down assay in hNPCs followed by western analysis shows that PARP1 and other SBBR core proteins preferentially bind wt AAV2 ITR compared to ΔBC ITR (n = 4 experiments; repeated measures one-way ANOVA of log_10_[protein expression] followed by Tukey’s test).

**Table 1.**
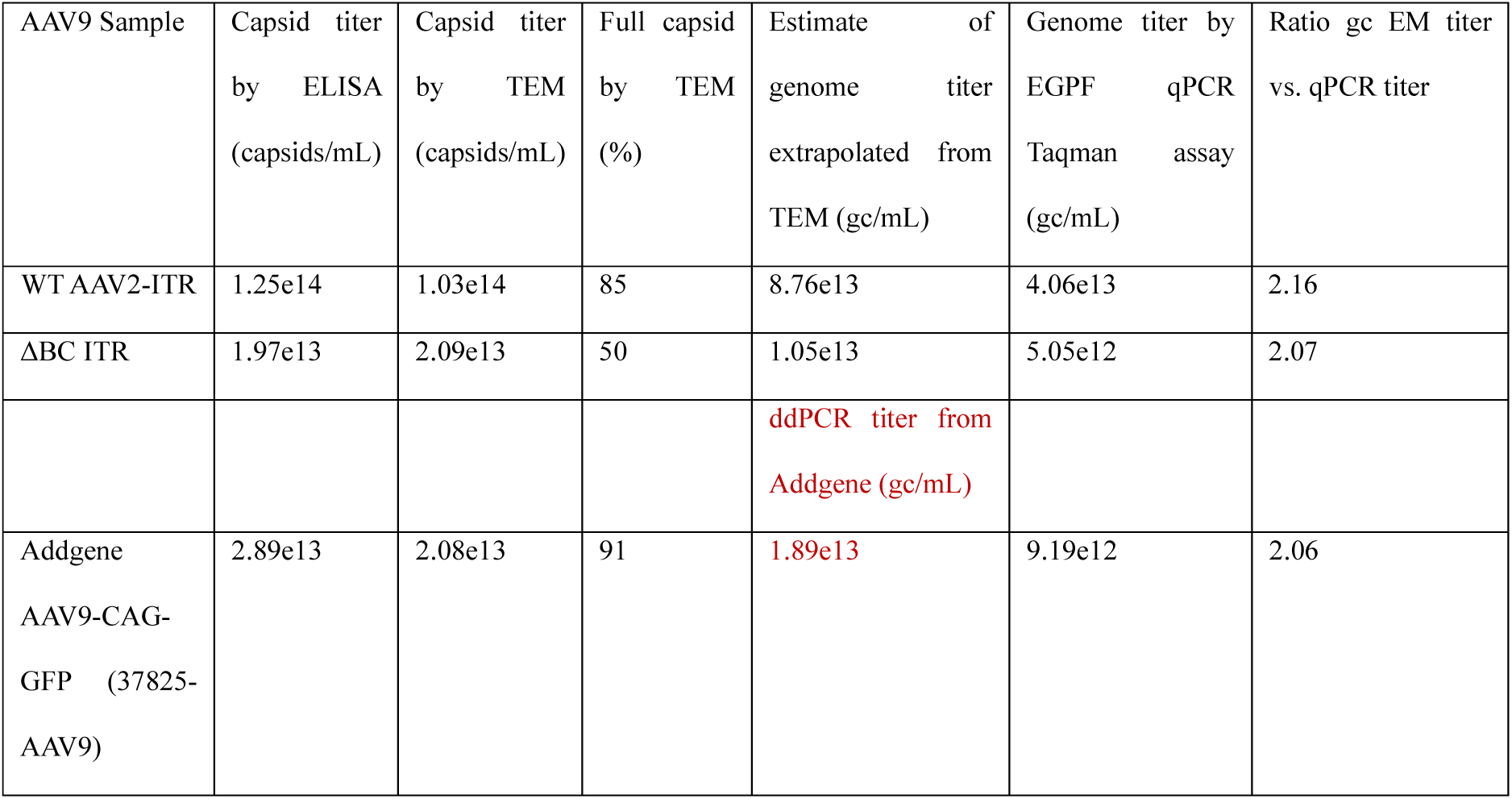
Multiple independent methods were used to estimate WT and ΔBC titer. AAV from Addgene (37825-AAV9) with known titer as measured by digital droplet PCR (ddPCR) by the supplier was used as control to estimate the titer of the custom preparations of AAV containing WT and ΔBC ITRs. First, the number of capsids were quantified by ELISA for each of the three samples. Next, an independent measurement of the number of total capsids in the Addgene preparation was estimated by assuming all full capsids contained complete genomes and dividing the ddPCR titer by the % of full capsids as measured by TEM. Then, capsid titers for the WT and ΔBC ITRs were calculated from TEM images by comparing the relative counts between each AAV and the Addgene AAV of known titer. This estimate of the capsid titer was in agreement with the capsid titers measured independently by ELISA. The genome titer was then estimated from the TEM images by multiplying the TEM capsid titers by the percentage of full capsids. Finally, this estimated genomic titer was compared to the titer obtained independently by qPCR for all three samples, which differed by approximately a factor of 2, consistent with known differences between quantification by ddPCR and qPCR.

Next, we tested to what extent the ΔBC ITR mutation impacts the toxicity of the virus. Figures 7C and 7D show data from time-lapse microscopy experiments comparing the toxicity of the ΔBC mutant to the wt control rAAV9 at various MOIs as measured by qPCR. As predicted, the impact of infection by the ΔBC mutant on hNPCs was significantly attenuated compared to the wt rAAV9 infection at the same MOI (reduction in proliferation at 1E6 gc/ml: wt 49 ± 3% vs ΔBC 25 ± 3%, p <0.0001; increase in cell death at 1E6 gc/ml: wt 120 ± 10% vs ΔBC 43 ± 5%, p <0.0001). Finally, we synthesized the ΔBC ITR DNA and compared its affinity for PARP1 and PARP1-associated proteins, identified from the affinity pull-down experiments described above (Fig. 3), to that of wt AAV2 ITR. Consistent with our hypothesis, wt AAV2 ITR bound PARP1 and other DDR proteins with higher affinity than the ΔBC ITR. Taken together, these data indicate that during rAAV infection, the T-shaped hairpin of the AAV ITR binds to and depletes host PARP1, analogous to pharmacologic PARP inhibitors, and that this interaction is necessary for inducing toxicity in dividing hNPCs.

## Discussion

### AAV ITRs binds host DDR proteins important for DNA replication, resulting in hNPC toxicity

We propose that during rAAV infection, AAV ITRs bind to and functionally deplete PARP1 and other DDR proteins, which induces replication stress and cell death in hNPCs. In support of this, we demonstrate that rAAV-toxicity mimics several key features of toxicity induced by the pharmacological PARP inhibitor ABT-888 in hNPCs. Similar to pharmacologic PARP inhibitors, rAAV is most toxic to dividing cells(*6*), with increasing rAAV toxicity observed with more rapid cell proliferation (Fig. 1). Both rAAV and ABT-888 inhibit PAR formation and induce a similar dose-dependent reduction in proliferation and increase in cell death (Figs. 1, 4, and 5). Moreover, the combined toxicity of low-dose rAAV and ABT-8880 is additive (Fig. 5), consistent with the predicted outcome of combining dose-dependent toxins that work through similar mechanisms. Our experiments also demonstrate that both agents induce aberrant cell cycle progression with nuclear enlargement and a similar pattern of activation of check point pathways (Fig. 2 and Fig. 6). In addition, both rAAV infection and ABT-888 treatment induce DNA double-strand breaks in the host genome (Fig. 3 and Fig. 6).

DNA damage can be induced in dividing cells, including NPCs, by inhibition of DNA polymerase and other enzymes important for DNA replication(*83*, *84*). Likewise, PARP1, other SSBR core proteins, and TOP1 serve several critical functions during mitosis, including participation in lagging strand synthesis and resolution of stalled DNA replication forks during S phase(*26–35*). There is a higher propensity for stalled replication forks to occur at repetitive DNA sequences, including repetitive GC-rich hairpin-forming DNA sequences like those found in AAV ITRs(*77–79*). The current results are also consistent with prior work demonstrating that DNA hairpins are among the DNA structures detected by PARP1(*62*, *63*). Thus, we propose that the T-shaped hairpin within AAV2 ITR is an excellent substrate for PARP1 binding, which depletes this important protein during cell division and contributes to rAAV-induced cellular toxicity. In support of this notion, mutating the T-shaped hairpins significantly reduces the binding of the AAV ITRs to PARP1 and other SSBR proteins and the toxicity of the virus in hNPCs.

### Recombinant AAV induces DNA damage

Previous studies demonstrate that infection with wildtype AAV induces a robust DNA damage response(*43–45*), but that infection with rAAV alone does not(*46*). Work by Hirsch et al. and Johnston et al. demonstrate that rAAV infection causes dose-dependent toxicity in dividing cells and that the AAV ITRs are likely sufficient and necessary for this effect(*6*, *19*). Building on these studies, the current experiments strongly suggest that ITRs recruit DDR proteins during rAAV infection and induce DNA damage in the host genome. These findings agree with two recent studies describing rAAV-induced expression of γH2AX+ and 53BP1+ foci in human induced pluripotent stem cell-derived neurons(*47*) and U2OS cells(*48*), respectively. While the reasons for the discrepancy with previous studies showing no DNA damage response by rAAV are likely multifactorial including differences in host cell type, it is important to note that rAAV toxicity (Fig. 1, Fig. 2C, Fig. 2G) and DNA damage (Fig. 3D) exhibits a steep dose dependence and could have been absent at the previously tested MOIs(*46*), which were lower than those used in this study.

### Engineering safer rAAV-based gene therapies

These dose-dependent effects described above also highlight the importance of carefully characterizing the therapeutic dose window for rAAV in the development of human gene therapies. This is particularly imperative in pediatric patients, who are still undergoing development and are the largest population to receive rAAV-based gene therapies. Extra caution should also be exercised when targeting adult organs, where proliferation of cells is ongoing and are critical for normal function. While there are reports of severe rAAV toxicity in postmitotic cells within the brain and other tissue (*1–4*, *7–11*), further work is needed to determine whether these toxicities are predominantly ITR-mediated and to what extent the same cellular pathways that are disrupted in dividing cells are also altered in postmitotic cells.

We present one of the first studies demonstrating that AAV ITRs can be engineered to reduce rAAV-induced toxicity(*20*). The ΔBC ITR mutant originally described by Zhou et al.(*81*) and investigated here retains a single hairpin, which is required to prime second-strand synthesis. The decreased toxicity exhibited by this mutant comes at a moderate cost of an 8-fold decrease in viral yield. In the future, the ΔBC rAAV or similar rAAV mutants engineered to contain reduced or altered hairpins in the ITRs could expand the therapeutic window of rAAV-based therapies, which is currently a fundamental obstacle to wider adoption of rAAV for human gene therapy(*10*, *85*).

### Harnessing the antiproliferative effects of rAAV for treating brain neoplasms

Previous work suggests that wildtype AAV could be effective as an oncolytic virus(*86*). However, its predilection to integrate into the host genome and replicate in the presence of a helper virus that commonly infects humans precludes its use as a viable therapeutic strategy. Other oncolytic viruses, such as herpes simplex, Zika virus, influenza, and adenovirus, have been investigated for the treatment of neoplasms in the brain and other organ systems. However, these viruses introduce a significant risk of systemic inflammation and encephalitis. In contrast, rAAV has been delivered intrathecally and intracranially in humans to treat genetic disease without evidence of widespread inflammation, although local cell loss around the injection site has been reported in some trials(*87*). The ability of AAV ITRs to inhibit PARP1 and other proteins important for cancer cell proliferation (Fig. S1) combined with their wide tropism, ability to accommodate DNA constructs up to ∼5,000 base pairs in length, and limited immunogenicity, make them attractive as candidates for engineering future oncolytic therapies. Once again, engineering of ITRs, in this case to make them more toxic to dividing cells, may provide additional therapeutic benefit.

### Inhibition of DDR proteins important for cell division may serve as a common mechanism among viruses that cause microcephaly

The complex biology of HSV, HIV, Zika, CMV, and Rubella viruses has thus far precluded our ability to identify a precise mechanism by which these infectious agents cause microcephaly. In contrast, rAAV, with its simple genome, inability to replicate, and lack of viral protein expression, offers a unique model system to probe mechanisms and potential therapies for viral-induced microcephaly that might be shared among these evolutionary distant viruses. Recent work by Rychlowska et al. demonstrates that Zika virus also induces DNA damage in the form of γH2AX activation and aberrant mitosis in human induced pluripotent stem cell-derived NPCs. The study demonstrates that Zika functionally inhibits PNKP, a SSBR core protein that binds AAV2 ITRs in our experiments (Fig. 3H and 3L). The nuclear entry of the Zika NS1 protein, rather than the Zika genome, triggers aberrant translocation of PNKP into the cytosol, depleting this enzyme from the nucleus. Unlike rAAV, Zika infection also inhibits rather than activates check points CHK1 and CHK2, resulting in mitotic catastrophe. Nonetheless, recruitment and resulting depletion of DDR proteins involved in DNA replication, including SSBR core proteins, in NPCs may be a common mechanism for viral-induced toxicity in microcephaly and other disorders(*88*). Many viruses, including those that cause microcephaly, have numerous hairpins in their genome that could be ideal binding substrates for DDR proteins(*89–92*). This might explain how distantly related viruses with no or little homology can impart similar toxicity in NPCs and induce microcephaly in humans. Future studies focused on shared mechanisms may lead to treatments for microcephaly and other neurodevelopmental disorders of viral etiology.

## Materials and Methods

### Study Design

The research objective was to investigates the cellular pathways and mechanisms underlying AAV toxicity in dividing cells that was previously identified by our laboratory(*6*) and others(*19*). No formal pre-specified hypotheses were outlined prior to the start of the study. Also, no power calculations were performed to estimate sample size and no primary or secondary endpoints were defined prior to the study. All treatments were randomized by well and control and experimental conditions were equally represented. Investigators performing imaging were blinded to sample identity. Investigators performing intermediate data analysis used automated software pipelines and were blinded to final outcome. Sample size varied depending on the experimental techniques being used. For cell proliferation (Incucyte) and histological measurements, experiments were repeated at least 3 times, with each experiment performed in triplicate (3 wells). Incucyte experiments described in Fig. 1 and Fig. 5 investigating the dependency of AAV and PARPi toxicity on treatment dose and rate of cell proliferation contained significantly more than 3 experiments. These were collected and analyzed retrospectively by pooling all Incucyte experiments testing AAV and PARPi toxicity containing all 3 treatment doses and saline control and exhibiting at least a 2-fold increase in proliferation under saline conditions. All western blot experiments were repeated 4 times. There were no data points excluded and no outliers were removed.

### Cell culture

H9 embryonic stem cells (WiCell) were differentiated into hNPCs as previously described(*93*). Briefly, H9 cells were cultured on Matrigel (Corning) coated plates with mTESR medium (Stemcell technologies) and passaged with collagenase IV (Gibco) upon reaching 80% confluence. To generate NPCs, H9 cells were transferred to ultra-low attachment cell culture plates (Corning) with mTESR medium + 10mM of Y-27632 (Tocris) in an orbital shaker for 24 hours, forming embryonic bodies (EBs). Subsequently, EBs were cultured in N2B27 media (Key Resources Table, Supplementary Information) using dual SMAD inhibition cocktail (500ng/ml Noggin; R&D Systems and 10mM SB431542; Tocris) for 10 days with media changes every other day. On day 10, EBs were plated on poly-ornithine (Sigma)/laminin (Life Technologies)-coated plates with N2B27 media + 1mg/ml laminin and allowed to attach for 24-36 hours. Finally, rosettes (neural precursor structures) were identified under the microscope and manually picked between 5-7 days, dissociated, and expanded as hNPCs. HNPCs were cultured in N2B27 media in the presence of FGF2 (20 ng/mL; Bio-Techne), using reduced growth factor basement membrane extract (Bio-Techne) coated plastic plates (Fisher Sci). Medium was changed every 2-3 days, and hNPCs were passaged with Accutase (Innovative Cell Technologies) when plates reached 90%-110% confluence. Experiments were only conducted with hNPCs with passage number ≤ 14.) U2OS cells (ATCC) were cultured in U2OS media (Key Resources Table, Supplementary Information) as previously described(*73*, *74*). U2OS cells stably expressing the PAR binding WWE domain linked to EGFP were generated (U2OS-LivePAR) as reported previously(*73–75*, *94*). CW468 patient-derived glioblastoma stem cells (GSCs) were previously isolated from a surgically resected tumor sample(*95*) and cultured in GSC media (Key Resources Table, Supplementary Information), similar to previously described methods(*96*). Cell cultures underwent regular testing for the presence of Mycoplasma using qPCR detection.

### In vitro rAAV transduction and time-lapse imaging

HNPCs or GSCs were counted using a hemocytometer (Fisher Sci) and seeded onto 96-well plates (Corning) at a density of 10k cells/well. After 18-24 hours, the medium was treated with propidium iodide (1 μg/mL; Sigma-Aldrich) along with rAAV (AAV1-CAG-flex-EGFP [Addgene], AAV9-ΔBC-ITR-CAG-GFP [Salk Viral Vector Core], or AAV9-WT-ITR-CAG-GFP [Salk Viral Vector Core] at MOI = 1E4, 1E5, or 1E6 or saline (equivalent volume to rAAV at MOI = 1E6) as previously described(*6*). For experiments testing PARP inhibitor (ABT-888; Sellechkem) or CHK2 inhibitor (BML-277; APExBIO), stock solutions were prepared in dimethyl sulfoxide (DMSO; Sigma-Aldrich) at 10 mM and 50 mM, respectively. Cells were treated with inhibitor or vehicle at the same timepoint as rAAV. Control wells contained the same amount of vehicle as used in experiments with the highest concentration of inhibitor. Nicotinamide riboside (NR) was purified from capsules (TruNiagen) by dissolving the contents in distilled water (Invitrogen) and filtering through a 0.2µm filter to remove the carrier. Five images per well were acquired 4 hours apart in bright field and red-fluorescence for 72 hours on an IncuCyte S3 Live Cell Analysis System (Sartorius) Each experiment was performed in triplicate, across at least three independent experiments. Data was extracted using IncuCyte Analysis software according to default analysis settings for phase and red channel. For each well, time-dependent confluence data were normalized by the confluence measurement at time = 0 to account for fluctuations in cell density and reported as a fold increase in confluence from time = 0. The percentage of cells labeled with PI for each well was measured by the total red fluorescence normalized by the (nonnormalized) confluence value at each timepoint. All statistical tests were performed using the last time point recorded at 72 hours. In rare instances where data were not collected or images were out of focus at 72 hours, the 68-hour time points were used for all groups in that experiment.

### Immunocytochemistry

Cells were seeded at a density of 10k/well into 18-well chamber slides (Ibidi). 18-24 hours later, cells were treated with experimental agents or corresponding vehicle controls at desired concentrations as described for time lapse imaging. After an additional 24 hours, cells were fixed in 4% formaldehyde prepared from powdered paraformaldehyde (Electron Microscopy Sciences) at room temperature for 45-60 minutes then rinsed with tris-buffered saline (TBS; Key Resources Table, Supplementary Information) three times for five minutes each. The samples were then incubated with blocking buffer (Key Resources Table, Supplementary Information) for 45-60 minutes at room temperature before overnight incubation with primary antibody at 4C. The following day, cells were rinsed twice for 5 minutes each with TBS before a 30-minute incubation with blocking buffer at room temperature. The cells were then incubated for two hours in the dark at room temperature with secondary antibody, rinsed with blocking buffer for 15 minutes, rinsed twice with TBS, incubated in 100 ng/ml DAPI (Sigma Aldrich) for 15 minutes, rinsed an additional two times with TBS, before one drop of mounting medium (Epredia) was placed into each well. Plates were imaged no sooner than 18 hours following application of mounting media using a Keyence BZ-X810 microscope using a 60X 1.4 oil immersion objective. Each experiment was repeated three times and performed in triplicate with five fields of view per well. Cellpose (version 3.1.1) was used for automated nuclear segmentation. Approximately 200 manually segmented nuclei from pilot experiments were used to fine-tune the Cellpose pretrained model (using default settings) across eight iterations, each consisting of one image(*97*). The average precision at the intersection over union (iou) threshold = 0.5 after the final iteration equaled 0.906. A custom Python script was used to calculate nuclear size from segmented nuclei. Nuclear puncta were segmented and quantified using Cell profiler’s pipeline for speckle counting (version 4.2.8) with empirical optimization or adjustment for puncta size and threshold. Primary antibodies and secondary antibodies used, along with the dilutions used for immunocytochemistry, can be found in the Key Resources Table, Supplementary Information.

### Western Blotting

300,000 cells/well were seeded into 6-well plates. 18-24 hours later cells were treated with AAV (AAV1-CAG-flex-EGFP) at an MOI of 1E5 or saline control. 18 hours post-treatment, cells were dissociated with Accutase, washed with PBS, and lysed with 1X RIPA Buffer Key Resources Table, Supplementary Information. Protein lysate supernatant was isolated from leftover cell debris using centrifugation at 14000 x g. Protein samples were collected and quantified by running a Bicinchoninic acid (BCA) assay according to manufacturer instructions (Thermo Scientific). Samples were prepared at 20ng protein/lane diluted in LDS Sample Buffer (Invitrogen). GAPDH (Cell Signaling Technology) was also stained and used as a positive control for each run and for normalization during quantitative western analysis. The samples were run on pre-cast gels (Invitrogen) and then transferred on to a Nitrocellulose (NC) membrane (Thermo Scientific) via a wet transfer process. Membrane was blocked in 3% BSA (Roche) in TBST (TBS and 0.1% Tween 20; Sigma Aldrich) or 5% non-fat milk (Research Products International) in TBST for one hour, depending on the antibody manufacturer’s recommendation. The membrane was then incubated in primary antibody overnight on a shaker in the cold room at 4C. The following day the membrane underwent washes with TBST and incubated in a secondary antibody for 1 hour at room temperature on a rocker. The blots were imaged using enhanced chemiluminescence western blotting detection reagents (Thermo Scientific) and imaged using the chemiluminescence channel on a Biorad ChemiDoc MP Imaging System. The primary and secondary antibodies used in these experiments can be found in Key Resources Table, Supplementary Information. In some cases, membranes were subsequently blocked for one hour and incubated with a new primary antibody overnight. In these cases, target proteins were separated in weight by at least one band on the protein ladder (Thermo Scientific) and exposure times for the second stain were at least tenfold lower than for the first stain, so that the two targets could be easily distinguished.

For quantification, the total intensity was plotted as a function of distance along each lane of the gel. Lane widths were set to be uniform and not overlapping. The background intensity was defined as the average of the five smallest intensity values identified in the area above each band, which typically contains less background staining than the area below the band. The background value of each lane was subtracted from the intensity plot for the corresponding lane. Any small negative values resulting from background subtraction were set to zero. The maximum value was determined for each lane and set as the center of the region of interest (ROI) used for quantification. In order to avoid subjective determination of the ROI, the superior border of the ROI was set to the location where the (background subtracted) intensity curve decayed to 5% of its maximum value. The inferior border was set to the same distance from the peak as the superior border such that the ROI was symmetric about the maximum. In cases where nonspecific staining was observed, a single peak matching the expected molecular weight from the literature was chosen. If another band was located above the band of interest, preventing complete decay of the intensity curve to 5% of its maximum value, the superior border of the ROI was set to the location where the intensity curve decayed to 5% above the local minimum between peaks. In cases where the control band was too faint to identify, superior and inferior borders for the experimental condition were used to delineate the ROI for the control condition. The background subtracted intensity curve was integrated between the borders of the ROI and normalized by the (background subtracted) intensity of the GAPDH band measured in the same sample. Different proteins measured in the same sample used the same GAPDH band for normalization.

### AAV ITR pulldown experiments

NPCs were collected from five to si× 10 cm dishes at 100%-120% confluence, washed twice with ice-cold DPBS, and resuspended in swelling buffer (Key Resources Table, Supplementary Information).Following 5 minutes of incubation in the swelling buffer, cells were manually scraped into 50 mL tubes. After centrifugation at 400 g for 10 minutes, the cell pellet was resuspended in lysis buffer(Key Resources Table, Supplementary Information), vortexed, and subjected to three additional centrifugation (600g) and washing steps in lysis buffer. Supernatant was discarded from the nuclei. For protein extraction, the nuclear pellet was resuspended in NP40 lysis buffer (Key Resources Table, Supplementary Information) with 1X protease/phosphatase inhibitor (ThermoFisher) and sonicated on ice at 30% power for 15 seconds followed by 45 seconds of rest. This was repeated 7 times with checks every two cycles to ensure that the mixture was still ice cold. Samples were subsequently rotated at 4C for 20 minutes, and centrifuged at 15,000 g at 4C for 20 minutes. The supernatant was pre-cleared using avidin beads for 30 minutes at room temp (Invitrogen), and the BCA assay was used for protein quantification. 10 µg of biotinylated DNA (IDT), which was annealed by heating to 95°C for 10 minutes and cooled slowly to room temperature, was incubated with 500 µg of pre-cleared protein extract for 1 hour at room temperature with rotation. Avidin beads (Invitrogen) were then added, and the mixture was incubated for an additional 2 hours at room temperature with rotation. Beads were washed 3x in NP40 lysis buffer and all supernatants were discarded. The beads were then incubated with protein loading buffer (LDS Sample Buffer 4X; Invitrogen) for denaturation at 100°C for 10 minutes and analyzed by western blot as described above (run at 20 μL of protein/lane without GAPD normalization) or by mass spectrometry.

Biotinylated DNA sequences (Integrated DNA Technologies; IDT) tested include wild-type AAV2, a scrambled ITR sequence controlling for GC content in the wild-type sequence, a random sequence that contains equal frequency of four nucleotides, and the ΔBC DNA sequence. The ΔBC sequence as originally described by Zhou et al. contains 111 bp. To account for length-dependent contribution to protein affinity, 34 base pairs were added from the 5’ end of the random sequence to the 5’ end of the ΔBC sequence to match the 145bp size of the AAV2 ITR, scrambled, and random oligomers. Secondary structure for each oligomer was modeled using the mFold algorithm through IDT’s OligoAnalyzer, using hairpin settings with SpecSheet Parameter Sets: Oligo Conc 0.25uM, Na+ Conc 50mM, Mg++ Conf 0mM, dNTPs Conc 0mM, and DNA Target Type). For each sequence, the most stable output was selected to represent the most likely secondary structure.

### RNAScope

Cells were seeded at a density of 10k/well into 18 well chamber slides (Ibidi). 18-24 hours later, cells were treated with experimental agents or corresponding vehicle controls at desired concentrations as described for time lapse imaging. After an additional 24 hours, RNAscope was performed using an probe for EGFP (ACD Bio) and Opal 520 fluorophore (Akoya Biosciences) to detect rAAV genomes according to the manufacturer’s Integrated Co-Detection Workflow (ACD Bio; Tech note MK 51-150)(*98*). This workflow was designed for use in tissue and required the following modifications: cells were prepared and pre-treated following Part 1 and Part 2, respectively, of the manufacturer’s Adherent Cell Protocol designed for 96 well plates (ACD Bio, SOP 45-009A_Tech Note)(*99*) before performing Part 4 of the Integrated Co-Detection Workflow. The protease digestion was also performed following the Adherent Cell Protocol rather than the Integrated Co-Detection. A detailed step by step protocol is available on the Shtrahman lab website and GitHub repository. Primary and secondary antibodies used for these experiments can be found in the Key Resources Table, Supplementary Information.

### Mass Spectrometry

Denatured protein samples were diluted in TNE buffer (Key Resources Table, Supplementary Information). RapiGest SF reagent (Waters Corp.) was added to the mix to a final concentration of 0.1% and samples were boiled for 5 min. TCEP (Tris (2-carboxyethyl) phosphine; ThermoFisher) was added to 1 mM (final concentration) and the samples were incubated at 37°C for 30 min. Subsequently, the samples were carboxymethylated with 0.5 mg/ml of iodoacetamide (ThermoFisher) for 30 min at 37°C followed by neutralization with 2 mM TCEP (final concentration). Proteins samples prepared as above were digested with trypsin (1:50 trypsin:protein ratio; ThermoFisher) overnight at 37°C. RapiGest was degraded and removed by treating the samples with 250 mM HCl (Fisher) at 37°C for 1 h followed by centrifugation at 14000 rpm for 30 min at 4°C. The soluble fraction was then added to a new tube and the peptides were extracted and desalted using C18 desalting columns (Thermo Scientific). Peptides were quantified using BCA assay (ThermoFisher) and a total of 1 µg of peptides were injected for LC-MS analysis.

Trypsin-digested peptides were analyzed by ultra high-pressure liquid chromatography (UPLC) coupled with tandem mass spectroscopy (LC-MS/MS) using nano-spray ionization. The nanospray ionization experiments were performed using an Orbitrap fusion Lumos hybrid mass spectrometer (Thermo) interfaced with nano-scale reversed-phase UPLC (Thermo Dionex UltiMate™ 3000 RSLC nano System) using a 25 cm, 75-micron ID glass capillary packed with 1.7-µm C18 (130) BEHTM beads (Waters Corporation). Peptides were eluted from the C18 column into the mass spectrometer using a linear gradient (5–80%) of Acetonitrile (ACN; Fisher) at a flow rate of 375 μl/min for 1h. The buffers used to create the ACN gradient were: Buffer A (Key Resources Table, Supplementary Information) and Buffer B (Key Resources Table, Supplementary Information). Mass spectrometer parameters are as follows; an MS1 survey scan using the orbitrap detector (mass range (m/z): 400-1500 (using quadrupole isolation), 120000 resolution setting, spray voltage of 2200 V, Ion transfer tube temperature of 275 C, AGC target of 400000, and maximum injection time of 50 ms) was followed by data dependent scans (top speed for most intense ions, with charge state set to only include +2-5 ions, and 5 second exclusion time, while selecting ions with minimal intensities of 50000 at which the collision event was carried out in the high energy collision cell (HCD Collision Energy of 30%), and the fragment masses were analyzed in the ion trap mass analyzer (With ion trap scan rate of turbo, first mass m/z was 100, AGC Target 5000 and maximum injection time of 35ms). Protein identification and label-free quantification were carried out using Peaks Studio 8.5 (Bioinformatics Solutions Inc.)

### Fluorescent Confocal Microscopy for the detection of poly(ADP-ribose) using LivePAR

Poly(ADP-ribose) (PAR) foci analysis was performed using U2OS cells expressing the genetically encoded LivePAR probe (U2OS/LivePAR), expressing a PAR binding domain (aa 100-182 from RNF146) fused to EGFP(*74*, *75*, *94*, *100*) .Microscope coverslips were washed sequentially in diethyl ether (Fisher Sci) for 20 minutes, followed by decreasing concentrations of ethanol (Fisher Sci), at 100%, 70%, and 0% in double-distilled water (ddH_2_O). The coverslips were then treated with 1 N HCl (Fisher Sci) for 20 minutes and then rinsed thrice with ddH_2_O. U2OS-LivePAR cells were exposed to AAV1-CAG-flex-EGFP (Addgene) for 8 hours before being transferred onto the prepared coverslips. After an additional 16 hours incubation, PARP-dependent PAR formation was induced by treating the cells with the methylating agent 1-methyl-3-nitro-1-nitrosoguanidine (10 µM, MNNG; Molport), the NAD+ precursor dihydronicotinamide riboside (100 µM, NRH; NuChem Sciences), and the PARG inhibitor PDD00017273 (10 µM; MilliporeSigma). To demonstrate PARP-dependent PAR formation, select cells were additionally treated with the PARP inhibitor ABT-888 (10 µM; Tocris) or DMSO (Fisher Sci) as a control. Following 90 minutes of exposure to non-viral treatments, cells were pre-fixed with 4% formaldehyde in PBS (Fisher Sci) for 15 minutes, subsequently fixed with ice-cold (−20°C) methanol (Fisher Sci):acetone (Fisher Sci) (7:3) for 9 minutes at −20°C, and stained using immunocytochemistry as previously described (*94*, *101*). The coverslips were washed with PBS (VWR), and blocked with 5% bovine serum albumin (BSA; RPI Research Products International) in 0.25% Triton X-100 (MilliporeSigma)/PBS (PBST) for one hour at room temperature. Following three washes with PBST, the coverslips were mounted onto glass slides using anti-fade mounting medium containing DAPI (Vectashield, Fisher Sci) and sealed with clear nail polish (O.P.I. Nature Strong). Images were acquired using a Nikon Ti2-E inverted confocal microscope with Ax-R equipped with Ax-R 2k Resonant + Galvo Scan Head. PAR foci per nucleus were quantified using ImageJ software (NIH).

### Flow Cytometry (FACS) Cell Cycle Analysis

HNPCs were plated at a density of 300,000 cells per well in 6-well plates. After 18 hours, cells were treated with AAV (AAV1-CAG-flex-EGFP; Addgene) at an MOI of 1e6 or saline. 24 hours post-infection, cells were collected using Accutase digestion and centrifuged at approximately 1000 × g for 6 minutes at room temperature. Supernatant was discarded and cell pellets were resuspended in 1 mL DPBS(-/-) (Cytiva, Corning), centrifuged again for 6 minutes at 200 x g and the supernatant was again discarded. Cell pellets were then resuspended in DPBS(-/-) to obtain a single cell suspension, which was kept on ice. The cell suspension was transferred slowly to tubes containing 1 mL of 70% ethanol (Sigma Aldrich), vortexed, and fixed for over 2 hours or overnight at 4°C. For DAPI staining, the fixed cells were centrifuged at 5000 × g for 15 minutes, ethanol was thoroughly removed, and pellets were resuspended in 1 mL DPBS(-/-) with 1% BSA, incubated for 1 minute, and centrifuged again at 5000 × g for 10 minutes. The cell pellets were then resuspended in 400 µL of DAPI/Triton X-100 staining solution and incubated in the dark at room temperature for 30 minutes.

The BD FACS Aria Fusion was used to acquire samples for cell cycle analysis. A 100 µm nozzle was used on the Aria Fusion, at a pressure of 20 psi. For cell cycle acquisition, the signal trigger was set for the DAPI channel with the signal threshold set to 5,000. Area scaling for the violet laser was adjusted to place the G0/G1 peak at a value of 50,000 in the DAPI Area parameter for each control sample. The histogram for some samples in the experimental group shows a rightward shift relative to the control due to cell loss and increased DAPI per cell. The data was then gated to include single cells, and these gated cells were subsequently analyzed for cell cycle using the FlowJo Cell Cycle function Watson (Pragmatic) model with default settings.

### Electron Microscopy Titer

Electron microscopy methods for imaging empty AAV capsids were adapted from previous protocols^6,98^. AAV9-WT-ITR-CAG-GFP (40x dilution) and AAV9-ΔBC-ITR-CAG-GFP (5x dilution) viruses (Salk Viral Vector Core), along with AAV9-CAG-GFP (20x dilution; Addgene), were submitted to the UCSD EM core for negative staining and Transmission Electron Microscopy (TEM) analysis to determine the empty-to-full capsid ratio. Dilutions were calculated by qPCR titer values for each virus to submit approximately 1e12gc/mL of each virus. 5 μL of each AAV dilution was applied for 5 minutes to glow-discharged formvar/carbon 400 mesh copper grids (Ted Pella). The grids were then washed three times with 0.22um filtered double-distilled water for 10 seconds and stained for 1 minute with 2% uranyl acetate (Ladd research) in double-distilled water. The grids were then blotted for 3 seconds onto Whatman grade 1 filter paper (Whatman) and air-dried. Images were acquired at 30,000x magnification using a Jeol 1400 plus transmission electron microscope operated at 80 KeV and equipped with a bottom-mounted 4kx4k Gatan one-view camera.

Manual counting of empty (donut shaped) and full (circular) capsids in TEM images was performed. The AAV9-CAG-GFP virus was found to contain 91% full capsids. The capsid titer was calculated by dividing the genomic (ddPCR) titer by the full ratio: (1.9e13 gc/mL) / 91% = 2.08e13 gc/mL. Next, by comparing the number of total capsids for the AAV9-WT-ITR-CAG-GFP and AAV9-ΔBC-ITR-CAG-GFP viruses to those of the Addgene virus obtained from TEM images at the same magnification and adjusting for dilution factors, we calculated the capsid titers for AAV9-WT-ITR-CAG-GFP and AAV9-ΔBC-ITR-CAG-GFP to be 1.03e14 capsids/mL and 2.09e13 capsids/mL, respectively. These capsid titers closely matched the results from an AAV9 capsid ELISA, which showed 1.25e14 capsids/mL for AAV9-WT-ITR-CAG-GFP and 1.97e13 capsids/mL for AAV9-ΔBC-ITR-CAG-GFP. Additionally, the TEM analysis revealed a full-to-empty ratio of 85% for AAV9-WT-ITR-CAG-GFP and 50% for AAV9-ΔBC-ITR-CAG-GFP. By multiplying the capsid titer by the full ratio, we estimated the genomic titers: AAV9-WT-ITR-CAG-GFP virus = 8.76e13 gc/mL and AAV9-ΔBC-ITR-CAG-GFP virus = 1.05e13 gc/mL. Notably, the gc titer determined by TEM counting was approximately two-fold higher than that determined by qPCR, consistent with the approximate two-three fold difference previously described between ddPCR and qPCR AAV titers(*82*).

### qPCR Viral Titer

QPCR TaqMan assay with EGFP-specific primers and probe were used to determine the viral genome titers for both WT-ITR and ΔBC-ITR AAVs. Two µL of viral sample was treated with DNase I (Thermo Scientific) in a 10 µL reaction at 37°C for 30 minutes, followed by inactivation using 1 µL of 50 mM EDTA at 65°C for 10 minutes. The DNase-treated sample was then subjected to alkaline digestion to release the AAV genomic DNA by incubating with 90 µL of alkaline digestion buffer (25 mM NaOH; Fisher BioReagents, 0.2 mM EDTA; EMD Millipore Corp) at 98°C for 8 minutes. The reaction was neutralized with 90 µL of neutralization buffer (40 mM Tris-HCl; Teknova pH 5, 0.05% Tween 20; Sigma Aldrich) on ice. The digested sample, containing the AAV genome, was diluted to fit within the standard curve range using ultrapure distilled water (Invitrogen) for the EGFP TaqMan assay. The qPCR was carried out using PrimeTime Gene Expression Master Mix (IDT) and the EGFP TaqMan Assay (ThermoFisher Scientific), with 2 µL of diluted AAV sample in a 20 µL reaction. The thermal cycling conditions were 95°C for 3 minutes, followed by 40 cycles of 95°C for 15 seconds and 60°C for 1 minute, performed on a Bio-Rad CFX96 Touch Real-Time PCR system. To quantify the viral genome copy numbers, a standard curve with eight-point 10-fold serial dilution (10^8^ – 10^1^ copies per reaction) was generated using KpnI-HF (NEB)-digested and linearized corresponding plasmids. All standards and samples were analyzed in technical triplicate, and no-template controls (NCs) included on every plate showed no amplification, with the standard-curve R^2^ > 0.99. As a control, we included AAV9-CAG-EGFP (Addgene) with a known titer determined by ddPCR to verify the EGFP TaqMan tittering assay.

### ELISA Viral Capsid Titer

Viral capsid titer was measured using an AAV9 Titration ELISA kit (Progen). Recombinant AAV samples were diluted to estimated concentrations within this manufactures recommended linear range (3e8–1.3e9 capsids/mL). The ELISA assay was carried out according to the manufacturer’s protocol and read using the Infinite 200Pro plate reader. Data analysis was performed using the MyAssays Online platform(*104*), applying a 5-parameter curve fit.

### Statistical analysis

Statistical comparisons were performed in Prism (GraphPad Software). Significance tests for experiments comparing AAV infection or PARP inhibitor treatment to control conditions were analyzed using unpaired two-tailed t-tests, except where described below. Significance tests for dose response data were analyzed using all to all one-way ANOVA followed by pairwise comparison using Tukey’s test. Statistical analysis of the number of PAR puncta under different conditions were performed using Kruskal-Wallis followed by Dunn’s multiple comparisons test for the following pairs of groups : (1) untreated vs. treated with MNNG, PARG inhibitor, and NRH (2) untreated vs. treated with AAV (3) untreated vs. treated with AAV, MNNG, PARG inhibitor, and NRH (4) treated with MNNG, PARG inhibitor, and NRH vs. treated with AAV, MNNG, PARG inhibitor, and NRH (5) treated with AAV vs. treated with AAV, MNNG, PARG inhibitor, and NRH (6) treated with MNNG, PARG inhibitor, and NRH vs. treated with PARP inhibitor, MNNG, PARG inhibitor, and NRH (7) treated with PARP inhibitor vs. treated with PARP inhibitor, MNNG, PARG inhibitor, and NRH. For statistical analysis of western blots comparing total protein ratios between different groups, either ratio paired t-tests of the intensity values (for experiments containing two groups) or repeated measures one-way ANOVA followed by pairwise comparison using Tukey’s test of the logarithm of the intensity values (for experiments containing three groups) were used. For time lapse imaging experiments, the best fit to an exponential plateau function y = a + b*exp(-k*x) or linear regression were calculated to describe the relationship between the rate of cell division and the reduction in proliferation or increase in cell death, respectively. Nonlinear Spearman (r_S_) or linear Pearson (r_P_) coefficients were calculated to quantify the correlation between the rate of cell division and the reduction in proliferation or increase in cell death, respectively. All data are presented as mean ± standard error of the mean. Error bars smaller than the symbols used to mark data points are not shown. The threshold for significance (α) was set at 0.05; n.s. is not significant, p< 0.05, 0.01, 0.001, 0.0001 are depicted respectively by *, **, ***, **** for all data.

## Acknowledgements

We thank Dr. Marianna Alperin, Dr. Jerome Mertens, Dr. Melissa Campbell, Dr. Catherine Freudenreich, and Brandon Jones for their comments on the manuscript and Ruth Keithley for her experimental advice regarding human neural progenitor cell culture. We would also like to acknowledge the UCSD Cellular and Molecular Medicine Electron Microscopy Core (UCSD-CMM-EM Core, RRID: SCR_022039) and the GT3 Viral Core Facility of the Salk Institute with funding from NIH-NCI (CCSG: P30 CA014195), for imaging and production of viral capsids, respectively. We are grateful to Cody Fine and Mitra Banihassan (UCSD) and the rest of the UCSD Human Embryonic Stem Cell Core Facility for technical assistance with flow cytometry and time lapse microscopy (Incucyte S3) experiments. The core is supported by the UCSD Stem Cell Program and a CIRM Major Facilities grant (FA1-00607) to the Sanford Consortium for Regenerative Medicine. The Incucyte S3 was purchased with funding from a National Institutes of Health grant (#1S10ODO25060-01).

## Author contributions

Data Curation: SF, MS, JZ, WPR

Formal Analysis: MS, SF, JG, C-HL, WPR, RA-R, DF, YH

Conceptualization: MS

Methodology: MS, RS, MR, MCM, JR, SF, JZ, NK, JG, C-HL, WPR, RA-R, FY, DF

Investigation: SF, JZ, GL, NK, JG, C-HL, WPR, RA-R, MW, FY, NL, JS, JL, EA-D, NS, ED

Funding acquisition: MS, RS

Project administration: MS, SF, NK, JG, C-HL, WPR, MW

Validation: SF, JZ, NK

Visualization: MS, SF, WPR

Supervision: MS, RS, MR, SF, JZ, NK, JG, MW

Resources: MS, RS, MCM, JR

Writing – original draft: MS, SF, NK, WPR

Writing – review & editing: MS, SF, RS, C-HL, WPR, MCM

## References

1. C. Hinderer, N. Katz, E. L. Buza, C. Dyer, T. Goode, P. Bell, L. K. Richman, J. M. Wilson, Severe Toxicity in Nonhuman Primates and Piglets Following High-Dose Intravenous Administration of an Adeno-Associated Virus Vector Expressing Human SMN. Hum. Gene Ther. 29, 285–298 (2018).

2. M. S. Keiser, P. T. Ranum, C. M. Yrigollen, E. M. Carrell, G. R. Smith, A. L. Muehlmatt, Y. H. Chen, J. M. Stein, R. L. Wolf, E. Radaelli, T. J. Lucas, P. Gonzalez-Alegre, B. L. Davidson, Toxicity after AAV delivery of RNAi expression constructs into nonhuman primate brain. Nat. Med. 27, 1982–1989 (2021).

3. A. Ulusoy, G. Sahin, T. Björklund, P. Aebischer, D. Kirik, Dose optimization for long-term rAAV-mediated RNA interference in the nigrostriatal projection neurons. Mol1. Ulusoy, A., Sahin, G., Björklund, T., Aebischer, P. Kirik, D. Dose Optim. long-term rAAV-mediated RNA Interf. nigrostriatal Proj. neurons. Mol. Ther. 17, 1574–1584 (2009).ecular Ther. 17, 1574–1584 (2009).

4. C. M. Suriano, J. L. Verpeut, N. Kumar, J. Ma, C. Jung, L. M. Boulanger, Adeno-associated virus (AAV) reduces cortical dendritic complexity in a TLR9-dependent manner. bioRxiv, 1–43 (2021).

5. Y. K. Chan, S. K. Wang, C. J. Chu, D. A. Copland, A. J. Letizia, H. C. Verdera, J. J. Chiang, M. Sethi, M. K. Wang, W. J. Neidermyer, Y. Chan, E. T. Lim, A. R. Graveline, M. Sanchez, R. F. Boyd, T. S. Vihtelic, R. G. C. O. Inciong, J. M. Slain, P. J. Alphonse, Y. Xue, L. R. Robinson-McCarthy, J. M. Tam, M. H. Jabbar, B. Sahu, J. F. Adeniran, M. Muhuri, P. W. L. Tai, J. Xie, T. B. Krause, A. Vernet, M. Pezone, R. Xiao, T. Liu, W. Wang, H. J. Kaplan, G. Gao, A. D. Dick, F. Mingozzi, M. A. McCall, C. L. Cepko, G. M. Church, Engineering adeno-associated viral vectors to evade innate immune and inflammatory responses. Sci. Transl. Med. 13, 1–18 (2021).

6. S. Johnston, S. L. Parylak, S. Kim, N. Mac, C. Lim, I. Gallina, C. Bloyd, A. Newberry, C. D. Saavedra, O. Novak, J. T. Gonçalves, F. H. Gage, M. Shtrahman, AAV ablates neurogenesis in the adult murine hippocampus. Elife. 10, 1–28 (2021).

7. P. Colella, G. Ronzitti, F. Mingozzi, Emerging Issues in AAV-Mediated In Vivo Gene Therapy. Mol. Ther. - Methods Clin. Dev. 8, 87–104 (2018).

8. M. Bugiani, T. E. M. Abbink, A. W. D. Edridge, L. van der Hoek, A. E. J. Hillen, N. P. van Til, G. V. Hu-A-Ng, M. Breur, K. Aiach, P. Drevot, M. Hocquemiller, R. Laufer, F. A. Wijburg, M. S. van der Knaap, Focal lesions following intracerebral gene therapy for mucopolysaccharidosis IIIA. Ann. Clin. Transl. Neurol., 904–917 (2023).

9. L. Servais, R. Horton, D. Saade, C. Bonnemann, F. Muntoni, D. O. Adjali, D. A. Beggs, D. D. Bharucha, D. C. Bönnemann, D. S. Braun, D. B. Byrne, D. M. Corti, D. A. Buj-Bello, D. J. Chamberlain, D. A. Ferreiro, D. K. Flanigan, M. O. Germanenko, D. N. Goemans, D. D. Grant, D. S. Hopkins, D. R. Horton, D. M. Kollb-Sielecka, D. C. Le Guiner, D. D. Levy, D. A. Lek, D. W. Miller, D. C. Morris, D. R. Dreghici, D. F. Muntoni, D. D. Saade, D. L. Servais, D. T. Singh, D. E. Vroom, D. K. Wagner, M. F. Van Ieperen, 261st ENMC International Workshop: Management of safety issues arising following AAV gene therapy. 17th-19th June 2022, Hoofddorp, The Netherlands. Neuromuscul. Disord. 33, 884–896 (2023).

10. S. Maurya, P. Sarangi, G. R. Jayandharan, Safety of Adeno-associated virus-based vector-mediated gene therapy—impact of vector dose. Cancer Gene Ther. 29, 1305–1306 (2022).

11. A. Kachanov, A. Kostyusheva, S. Brezgin, I. Karandashov, N. Ponomareva, A. Tikhonov, A. Lukashev, V. Pokrovsky, A. A. Zamyatnin, A. Parodi, V. Chulanov, D. Kostyushev, The menace of severe adverse events and deaths associated with viral gene therapy and its potential solution. Med. Res. Rev. 44, 2112–2193 (2024).

12. A. Philippidis, Novartis Confirms Deaths of Two Patients Treated with Gene Therapy Zolgensma. Hum. Gene Ther. 33, 842–844 (2022).

13. J. Guillou, A. de Pellegars, F. Porcheret, V. Frémeaux-Bacchi, E. Allain-Launay, C. Debord, M. Denis, Y. Péréon, C. Barnérias, I. Desguerre, G. Roussey, S. Mercier, Fatal thrombotic microangiopathy case following adeno-associated viral SMN gene therapy. Blood Adv. 6, 4266–4270 (2022).

14. A. Philippidis, Fourth Boy Dies in Clinical Trial of Astellas’ AT132. Hum. Gene Ther. 32, 1008–1010 (2021).

15. A. Lek, B. Wong, A. Keeler, M. Blackwood, K. Ma, S. Huang, K. Sylvia, A. R. Batista, R. Artinian, D. Kokoski, S. Parajuli, J. Putra, C. K. Carreon, H. Lidov, K. Woodman, S. Pajusalu, J. M. Spinazzola, T. Gallagher, J. LaRovere, D. Balderson, L. Black, K. Sutton, R. Horgan, M. Lek, T. Flotte, Death after High-Dose rAAV9 Gene Therapy in a Patient with Duchenne’s Muscular Dystrophy. N. Engl. J. Med. 389, 1203–1210 (2023).

16. A. Philippidis, After Patient Death, FDA Places Hold on Pfizer Duchenne Muscular Dystrophy Gene Therapy Trial. Hum. Gene Ther. 33, 111–115 (2022).

17. D. Duan, Lethal immunotoxicity in high-dose systemic AAV therapy. Mol. Ther. 31, 3123–3126 (2023).

18. M. A. Kotterman, D. V. Schaffer, Engineering adeno-associated viruses for clinical gene therapy. Nat. Rev. Genet. 15, 445–451 (2014).

19. M. L. Hirsch, B. M. Fagan, R. Dumitru, J. J. Bower, S. Yadav, M. H. Porteus, L. H. Pevny, R. J. Samulski, Viral Single-Strand DNA Induces p53-Dependent Apoptosis in Human Embryonic Stem Cells. 6, 1–13 (2011).

20. L. Song, T. Hasegawa, N. J. Brown, J. J. Bower, R. J. Samulski, M. L. Hirsch, AAV vector transduction restriction and attenuated toxicity in hESCs via a rationally designed inverted terminal repeat. Nucleic Acids Res. (2025).

21. P. P. Garcez, E. C. Loiola, R. Madeiro da Costa, L. M. Higa, P. Trindade, R. Delvecchio, J. M. Nascimento, R. Brindeiro, A. Tanuri, S. K. Rehen, Zika virus impairs growth in human neurospheres and brain organoids. Science. 352, 816–8 (2016).

22. D. M. P. Lawrence, L. C. Durham, L. Schwartz, P. Seth, D. Maric, E. O. Major, Human immunodeficiency virus type 1 infection of human brain-derived progenitor cells. J. Virol. 78, 7319–28 (2004).

23. L. Schwartz, L. Civitello, A. Dunn-Pirio, S. Ryschkewitsch, E. Berry, W. Cavert, N. Kinzel, D. M. P. Lawrence, R. Hazra, E. O. Major, Evidence of human immunodeficiency virus type 1 infection of nestin-positive neural progenitors in archival pediatric brain tissue. J. Neurovirol. 13, 274–283 (2007).

24. M. Rolland, X. Li, Y. Sellier, H. Martin, T. Perez-Berezo, B. Rauwel, A. Benchoua, B. Bessières, J. Aziza, N. Cenac, M. Luo, C. Casper, M. Peschanski, D. Gonzalez-Dunia, M. Leruez-Ville, C. Davrinche, S. Chavanas, PPARγ Is Activated during Congenital Cytomegalovirus Infection and Inhibits Neuronogenesis from Human Neural Stem Cells. PLOS Pathog. 12, e1005547 (2016).

25. M. H. Luo, H. Hannemann, A. S. Kulkarni, P. H. Schwartz, J. M. O’Dowd, E. A. Fortunato, Human cytomegalovirus infection causes premature and abnormal differentiation of human neural progenitor cells. J. Virol. 84, 3528–41 (2010).

26. D. Slade, Mitotic functions of poly(ADP-ribose) polymerases. Biochem. Pharmacol. 167 (2019), pp. 33–43.

27. G. E. Ronson, A. L. Piberger, M. R. Higgs, A. L. Olsen, G. S. Stewart, P. J. McHugh, E. Petermann, N. D. Lakin, PARP1 and PARP2 stabilise replication forks at base excision repair intermediates through Fbh1-dependent Rad51 regulation. Nat. Commun. 9 (2018), doi:10.1038/s41467-018-03159-2.

28. W. Min, C. Bruhn, P. Grigaravicius, Z. W. Zhou, F. Li, A. Krüger, B. Siddeek, K. O. Greulich, O. Popp, C. Meisezahl, C. F. Calkhoven, A. Bürkle, X. Xu, Z. Q. Wang, Poly(ADP-ribose) binding to Chk1 at stalled replication forks is required for S-phase checkpoint activation. Nat. Commun. 4 (2013), doi:10.1038/ncomms3993.

29. H. E. Bryant, E. Petermann, N. Schultz, A. S. Jemth, O. Loseva, N. Issaeva, F. Johansson, S. Fernandez, P. McGlynn, T. Helleday, PARP is activated at stalled forks to mediate Mre11-dependent replication restart and recombination. EMBO J. 28, 2601–2615 (2009).

30. A. Kaima Tsukada, T. Miyake, R. Imamura, K. Saikawa, M. Saito, N. Kase, M. Ishiai, Y. Matsumoto, M. Shimada, CDKs-mediated phosphorylation of PNKP is required for end-processing of single-strand DNA gaps on Okazaki Fragments and genome stability Short title: Phosphorylation of PNKP by CDK for processing single-strand DNA gaps (2024), doi:10.1101/2021.07.29.452278.

31. H. Hanzlikova, I. Kalasova, A. A. Demin, L. E. Pennicott, Z. Cihlarova, K. W. Caldecott, The Importance of Poly(ADP-Ribose) Polymerase as a Sensor of Unligated Okazaki Fragments during DNA Replication. Mol. Cell. 71, 319–331.e3 (2018).

32. H. Sun, L. Ma, Y. F. Tsai, T. Abeywardana, B. Shen, L. Zheng, Okazaki fragment maturation: DNA flap dynamics for cell proliferation and survival. Trends Cell Biol. 33 (2023), pp. 221–234.

33. S. Kumamoto, A. Nishiyama, Y. Chiba, R. Miyashita, C. Konishi, Y. Azuma, M. Nakanishi, HPF1-dependent PARP activation promotes LIG3-XRCC1-mediated backup pathway of Okazaki fragment ligation. Nucleic Acids Res. 49, 5003–5016 (2021).

34. A. Vaitsiankova, K. Burdova, M. Sobol, A. Gautam, O. Benada, H. Hanzlikova, K. W. Caldecott, PARP inhibition impedes the maturation of nascent DNA strands during DNA replication. Nat. Struct. Mol. Biol. 29, 329–338 (2022).

35. S. P. Chowdhuri, B. B. Das, Top1-PARP1 association and beyond: From DNA topology to break repair. NAR Cancer. 3 (2021), doi:10.1093/narcan/zcab003.

36. C. Hammack, S. C. Ogden, J. C. Madden, A. Medina, C. Xu, E. Phillips, Y. Son, A. Cone, S. Giovinazzi, R. A. Didier, D. M. Gilbert, H. Song, G. Ming, Z. Wen, M. A. Brinton, A. Gunjan, H. Tang, J. Virol., in press, doi:10.1128/JVI.00638-19.

37. M. Rychlowska, A. Agyapong, M. Weinfeld, L. M. Schang, Zika Virus Induces Mitotic Catastrophe in Human Neural Progenitors by Triggering Unscheduled Mitotic Entry in the Presence of DNA Damage While Functionally Depleting Nuclear PNKP. J. Virol. 96 (2022), doi:10.1128/jvi.00333-22.

38. K. Maeshima, H. Iino, S. Hihara, N. Imamoto, Nuclear size, nuclear pore number and cell cycle. Nucleus. 2, 113–118 (2011).

39. H. B. Steen, T. Lindmo, Cellular and nuclear volume during the cell cycle of NHIK 3025 cells. Cell Tissue Kinet. 11, 69–81 (1978).

40. J. Fidorra, T. Mielke, J. Booz, L. E. Feinendegen, Cellular and nuclear volume of human cells during the cell cycle. Radiat. Environ. Biophys. 19, 205–214 (1981).

41. J. G. Umen, The elusive sizer. Curr. Opin. Cell Biol. 17, 435–441 (2005).

42. S. Lodovichi, T. Cervelli, A. Pellicioli, A. Galli, Inhibition of dna repair in cancer therapy: Toward a multi-target approach. Int. J. Mol. Sci. 21, 1–29 (2020).

43. R. A. Schwartz, C. T. Carson, C. Schuberth, M. D. Weitzman, Adeno-Associated Virus Replication Induces a DNA Damage Response Coordinated by DNA-Dependent Protein Kinase. J. Virol. 83, 6269–6278 (2009).

44. J. Jurvansuu, K. Raj, A. Stasiak, P. Beard, Viral Transport of DNA Damage That Mimics a Stalled Replication Fork. J. Virol. 79, 569–580 (2005).

45. K. Ning, C. A. Kuz, F. Cheng, Z. Feng, Z. Yan, J. Qiu, Adeno-Associated Virus Monoinfection Induces a DNA Damage Response and DNA Repair That Contributes to Viral DNA Replication. MBio. 14 (2023), doi:10.1128/mbio.03528-22.

46. M. Fragkos, M. Breuleux, N. Clément, P. Beard, Recombinant Adeno-Associated Viral Vectors Are Deficient in Provoking a DNA Damage Response. J. Virol. 82, 7379–7387 (2008).

47. H. Costa-Verdera, O. San Raffaele, V. Meneghini, Z. Fitzpatrick, M. A. Alezz, E. Fabyanic, X. Huang, Y. Dzhashiashvili, A. Ahiya, E. Mangiameli, E. Valeri, G. Crivicich, I. Cuccovillo, R. Caccia, B. Bertin, G. Ronzitti, E. Engel, I. Merelli, F. Mingozzi, A. Gritti, K. Kuranda, A. Kajaste-Rudnitski, AAV vectors trigger DNA damage responses and STING-dependent inflammation in human CNS cells. Nat. Portf. Prepr., 1–34 (2024).

48. A. C. Maurer, B. Benyamini, V. B. Fan, O. N. Whitney, G. M. Dailey, X. Darzacq, M. D. Weitzman, R. Tjian, Double-Strand Break Repair Pathways Differentially Affect Processing and Transduction Double-Strand Break Repair Pathways Differentially Affect Processing and Transduction 2 by Dual AAV Vectorsby Dual AAV Vectors. bioRxiviv. (2023), 10.1101/2023.09.19.558438.

49. V. Turinetto, C. Giachino, Multiple facets of histone variant H2AX: A DNA double-strand-break marker with several biological functions. Nucleic Acids Res. 43, 2489–2498 (2015).

50. A. Shibata, P. A. Jeggo, Roles for 53BP1 in the repair of radiation-induced DNA double strand breaks. DNA Repair (Amst). 93, 102915 (2020).

51. K. H. Almeida, R. W. Sobol, A unified view of base excision repair: Lesion-dependent protein complexes regulated by post-translational modification. DNA Repair (Amst). 6, 695–711 (2007).

52. R. Abbotts, D. M. Wilson, Coordination of DNA single strand break repair. Free Radic. Biol. Med. 107, 228–244 (2017).

53. K. W. Caldecott, Causes and consequences of DNA single-strand breaks. Trends Biochem. Sci. 49, 68–78 (2024).

54. S. P. Chowdhuri, B. B. Das, Top1-PARP1 association and beyond: From DNA topology to break repair. NAR Cancer. 3, 1–8 (2021).

55. S. K. Das, I. Rehman, A. Ghosh, S. Sengupta, P. Majumdar, B. Jana, B. B. Das, Poly(ADP-ribose) polymers regulate DNA topoisomerase i (Top1) nuclear dynamics and camptothecin sensitivity in living cells. Nucleic Acids Res. 44, 8363–8375 (2016).

56. R. Hailstone, R. Maroofian, L. Woodbine, E. Korneeva, J. Brazina, A. Macaya, M. Severino, H. Tomoum, H. Houlden, K. W. Caldecott, medRxiv, in press (available at http://medrxiv.org/content/early/2023/06/09/2023.06.09.23291078.abstract).

57. M. R. Garrelfs, S. Takada, E. J. Kamsteeg, S. Pegge, G. Mancini, M. Engelen, B. van de Warrenburg, A. Rennings, J. van Gaalen, I. Peters, C. Weemaes, M. van der Burg, M. A. Willemsen, The Phenotypic Spectrum of PNKP-Associated Disease and the Absence of Immunodeficiency and Cancer Predisposition in a Dutch Cohort. Pediatr. Neurol. 113, 26–32 (2020).

58. G. Fragola, A. M. Mabb, B. Taylor-Blake, J. K. Niehaus, W. D. Chronister, H. Mao, J. M. Simon, H. Yuan, Z. Li, M. J. McConnell, M. J. Zylka, Deletion of Topoisomerase 1 in excitatory neurons causes genomic instability and early onset neurodegeneration. Nat. Commun. 11 (2020), doi:10.1038/s41467-020-15794-9.

59. N. C. Hoch, H. Hanzlikova, S. L. Rulten, M. Tétreault, E. Komulainen, L. Ju, P. Hornyak, Z. Zeng, W. Gittens, S. A. Rey, K. Staras, G. M. S. Mancini, P. J. McKinnon, Z. Q. Wang, J. D. Wagner, G. Yoon, K. W. Caldecott, XRCC1 mutation is associated with PARP1 hyperactivation and cerebellar ataxia. Nature. 541, 87–91 (2017).

60. E. O’Connor, J. Vandrovcova, E. Bugiardini, V. Chelban, A. Manole, I. Davagnanam, S. Wiethoff, A. Pittman, D. S. Lynch, S. Efthymiou, S. Marino, A. Y. Manzur, M. Roberts, M. G. Hanna, H. Houlden, E. Matthews, N. W. Wood, Mutations in XRCC1 cause cerebellar ataxia and peripheral neuropathy. J. Neurol. Neurosurg. Psychiatry. 0, 2017–2019 (2018).

61. R. Owczarzy, A. V. Tataurov, Y. Wu, J. A. Manthey, K. A. McQuisten, H. G. Almabrazi, K. F. Pedersen, Y. Lin, J. Garretson, N. O. McEntaggart, C. A. Sailor, R. B. Dawson, A. S. Peek, IDT SciTools: a suite for analysis and design of nucleic acid oligomers. Nucleic Acids Res. 36 (2008), doi:10.1093/nar/gkn198.

62. N. Laspata, D. Muoio, E. Fouquerel, Multifaceted Role of PARP1 in Maintaining Genome Stability Through Its Binding to Alternative DNA Structures. J. Mol. Biol. 436, 168207 (2024).

63. N. V. Malyuchenko, E. Y. Kotova, M. P. Kirpichnikov, V. M. Studitsky, A. V. Feofanov, PARP1 Binding to DNA Breaks and Hairpins Alters Nucleosome Structure. Moscow Univ. Biol. Sci. Bull. 74, 158–162 (2019).

64. A. Wilk, F. Hayat, R. Cunningham, J. Li, S. Garavaglia, L. Zamani, D. M. Ferraris, P. Sykora, J. Andrews, J. Clark, A. Davis, L. Chaloin, M. Rizzi, M. Migaud, R. W. Sobol, Extracellular NAD+ enhances PARP-dependent DNA repair capacity independently of CD73 activity. Sci. Rep. 10, 1–21 (2020).

65. M. M. Murata, X. Kong, E. Moncada, Y. Chen, H. Imamura, P. Wang, M. W. Berns, K. Yokomori, M. A. Digman, NAD+ consumption by PARP1 in response to DNA damage triggers metabolic shift critical for damaged cell survival. Mol. Biol. Cell. 30, 2584–2597 (2019).

66. C. C. Alano, P. Garnier, W. Ying, Y. Higashi, T. M. Kauppinen, R. A. Swanson, NAD+ depletion is necessary and sufficient for poly(ADP-ribose) polymerase-1-mediated neuronal death. J. Neurosci. 30, 2967–2978 (2010).

67. E. Fouquerel, E. M. Goellner, Z. Yu, J. P. Gagné, M. B. de Moura, T. Feinstein, D. Wheeler, P. Redpath, J. Li, G. Romero, M. Migaud, B. Van Houten, G. G. Poirier, R. W. Sobol, ARTD1/PARP1 negatively regulates glycolysis by inhibiting hexokinase 1 independent of NAD+ depletion. Cell Rep. 8, 1819–1831 (2014).

68. Y. Hou, S. Lautrup, S. Cordonnier, Y. Wang, D. L. Croteau, E. Zavala, Y. Zhang, K. Moritoh, J. F. O’Connell, B. A. Baptiste, T. V. Stevnsner, M. P. Mattson, V. A. Bohr, NAD+ supplementation normalizes key Alzheimer’s features and DNA damage responses in a new AD mouse model with introduced DNA repair deficiency. Proc. Natl. Acad. Sci. U. S. A. 115, E1876–E1885 (2018).

69. F. Salech, D. P. Ponce, A. C. Paula-Lima, C. D. SanMartin, M. I. Behrens, Nicotinamide, a Poly [ADP-Ribose] Polymerase 1 (PARP-1) Inhibitor, as an Adjunctive Therapy for the Treatment of Alzheimer’s Disease. Front. Aging Neurosci. 12, 1–10 (2020).

70. A. R. Lehmann, S. Kirk-Bell, S. Shall, W. J. D. Whish, “THE RELATIONSHIP BETWEEN CELL GROWTH, MACROMOLECULAR SYNTHESIS AND POLY ADP-RIBOSE POLYMERASE IN LYMPHOID CELLS” (1974).

71. J. Pietrzak, C. M. Spickett, T. Płoszaj, L. Virág, A. Robaszkiewicz, PARP1 promoter links cell cycle progression with adaptation to oxidative environment. Redox Biol. 18 (2018), pp. 1–5.

72. M. Rose, J. T. Burgess, K. O’Byrne, D. J. Richard, E. Bolderson, PARP Inhibitors: Clinical Relevance, Mechanisms of Action and Tumor Resistance. Front. Cell Dev. Biol. 8, 1–22 (2020).

73. C. A. Koczor, K. M. Saville, J. F. Andrews, J. Clark, Q. Fang, J. Li, R. Q. Al-Rahahleh, M. Ibrahim, S. McClellan, M. V. Makarov, M. E. Migaud, R. W. Sobol, Temporal dynamics of base excision/single-strand break repair protein complex assembly/disassembly are modulated by the PARP/NAD+/SIRT6 axis. Cell Rep. 37 (2021), doi:10.1016/j.celrep.2021.109917.

74. C. A. Koczor, A. J. Haider, K. M. Saville, J. Li, J. F. Andrews, A. V. Beiser, R. W. Sobol, Live Cell Detection of Poly(ADP-Ribose) for Use in Genetic and Genotoxic Compound Screens. Cancers (Basel). 14 (2022), doi:10.3390/cancers14153676.

75. C. A. Koczor, K. M. Saville, R. Q. Al-Rahahleh, J. F. Andrews, J. Li, R. W. Sobol, in Methods in Molecular Biology (Humana Press Inc., 2023), vol. 2609, pp. 43–59.

76. J. Li, K. M. Saville, M. Ibrahim, X. Zeng, S. McClellan, A. Angajala, A. Beiser, J. F. Andrews, M. Sun, C. A. Koczor, J. Clark, F. Hayat, M. V. Makarov, A. Wilk, N. A. Yates, M. E. Migaud, R. W. Sobol, NAD+bioavailability mediates PARG inhibition-induced replication arrest, intra S-phase checkpoint and apoptosis in glioma stem cells. NAR Cancer. 3 (2021), doi:10.1093/narcan/zcab044.

77. I. Voineagu, V. Narayanan, K. S. Lobachev, S. M. Mirkin, “Replication stalling at unstable inverted repeats: Interplay between DNA hairpins and fork stabilizing proteins” (2008).

78. S. Lu, G. Wang, A. Bacolla, J. Zhao, S. Spitser, K. M. Vasquez, Short inverted repeats are hotspots for genetic instability: Relevance to cancer genomes. Cell Rep. 10, 1674–1680 (2015).

79. S. Kaushal, C. H. Freudenreich, The role of fork stalling and DNA structures in causing chromosome fragility. Genes Chromosom. Cancer. 58, 270–283 (2019).

80. D. Slade, PARP and PARG inhibitors in cancer treatment. Genes Dev. 34, 360–394 (2020).

81. Q. Zhou, W. Tian, C. Liu, Z. Lian, X. Dong, X. Wu, Deletion of the B-B’ and C-C’ regions of inverted terminal repeats reduces rAAV productivity but increases transgene expression. Sci. Rep. 7, 1–13 (2017).

82. Addgene, Viral Production, (available at https://www.addgene.org/viral-service/viral-production/).

83. N. Michel, H. M. R. Young, N. D. Atkin, U. Arshad, R. Al-Humadi, S. Singh, A. Manukyan, L. Gore, I. E. Burbulis, Y. H. Wang, M. J. McConnell, Transcription-associated DNA DSBs activate p53 during hiPSC-based neurogenesis. Sci. Rep. 12, 1–14 (2022).

84. M. Wang, P. C. Wei, C. K. Lim, I. S. Gallina, S. Marshall, M. C. Marchetto, F. W. Alt, F. H. Gage, Increased Neural Progenitor Proliferation in a hiPSC Model of Autism Induces Replication Stress-Associated Genome Instability. Cell Stem Cell. 26, 221–233.e6 (2020).

85. N. Paulk, Gene Therapy: It Is Time to Talk about High-Dose AAV. Genet. Eng. Biotechnol. News. 40, 14–16 (2020).

86. K. Raj, P. Ogston, P. Beard, Virus-mediated killing of cells that lack p53 activity. Nature. 412, 914–917 (2001).

87. A. Philippidis, “‘Profoundly saddened’” Lysogene discloses child’s death in phase II/III trial. Hum. Gene Ther. 31, 1141–1143 (2020).

88. L. N. Lupey-Green, S. A. Moquin, K. A. Martin, S. M. McDevitt, M. Hulse, L. B. Caruso, R. T. Pomerantz, J. L. Miranda, I. Tempera, PARP1 restricts Epstein Barr Virus lytic reactivation by binding the BZLF1 promoter. Virology. 507, 220–230 (2017).

89. R. G. Huber, X. N. Lim, W. C. Ng, A. Y. L. Sim, H. X. Poh, Y. Shen, S. Y. Lim, K. B. Sundstrom, X. Sun, J. G. Aw, H. K. Too, P. H. Boey, A. Wilm, T. Chawla, M. M. Choy, L. Jiang, P. F. de Sessions, X. J. Loh, S. Alonso, M. Hibberd, N. Nagarajan, E. E. Ooi, P. J. Bond, O. M. Sessions, Y. Wan, Structure mapping of dengue and Zika viruses reveals functional long-range interactions. Nat. Commun. 10 (2019), doi:10.1038/s41467-019-09391-8.

90. A. J. Rennekamp, P. Wang, P. M. Lieberman, Evidence for DNA Hairpin Recognition by Zta at the Epstein-Barr Virus Origin of Lytic Replication. J. Virol. 84, 7073–7082 (2010).

91. H. L. Nakhasi, X. Q. Cao, T. A. Rouault, T. Y. Liu, Specific binding of host cell proteins to the 3’-terminal stem-loop structure of rubella virus negative-strand RNA. J. Virol. 65, 5961–5967 (1991).

92. S. K. Weller, D. M. Coen, Herpes simplex viruses: Mechanisms of DNA replication. Cold Spring Harb. Perspect. Biol. 4 (2012), doi:10.1101/cshperspect.a013011.

93. A. Sarkar, A. Mei, A. C. M. Paquola, S. Stern, C. Bardy, J. R. Klug, S. Kim, N. Neshat, H. J. Kim, M. Ku, M. N. Shokhirev, D. H. Adamowicz, M. C. Marchetto, R. Jappelli, J. A. Erwin, K. Padmanabhan, M. Shtrahman, X. Jin, F. H. Gage, Efficient Generation of CA3 Neurons from Human Pluripotent Stem Cells Enables Modeling of Hippocampal Connectivity In Vitro. Cell Stem Cell. 22, 684–697.e9 (2018).

94. R. Q. Al-Rahahleh, W. P. Roos, K. M. Saville, J. F. Andrews, Z. Wu, C. A. Koczor, A. Prakash, R. W. Sobol, Overexpression of the WWE domain of RNF146 modulates poly-(ADP)-ribose dynamics at sites of DNA damage. DNA Repair (Amst). 150, 103845 (2025).

95. D. Dixit, B. C. Prager, R. C. Gimple, H. X. Poh, Y. Wang, Q. Wu, Z. Qiu, R. L. Kidwell, L. J. Y. Kim, Q. Xie, K. Vitting-Seerup, S. Bhargava, Z. Dong, L. Jiang, Z. Zhu, P. Hamerlik, S. R. Jaffrey, J. C. Zhao, X. Wang, J. N. Rich, The RNA m6A Reader YTHDF2 Maintains Oncogene Expression and Is a Targetable Dependency in Glioblastoma Stem Cells. Cancer Discov. 11, 480–499 (2021).

96. M. Tang, Q. Xie, R. C. Gimple, Z. Zhong, T. Tam, J. Tian, R. L. Kidwell, Q. Wu, B. C. Prager, Z. Qiu, A. Yu, Z. Zhu, P. Mesci, H. Jing, J. Schimelman, P. Wang, D. Lee, M. H. Lorenzini, D. Dixit, L. Zhao, S. Bhargava, T. E. Miller, X. Wan, J. Tang, B. Sun, B. F. Cravatt, A. R. Muotri, S. Chen, J. N. Rich, Three-dimensional bioprinted glioblastoma microenvironments model cellular dependencies and immune interactions. Cell Res. 30, 833–853 (2020).

97. C. Stringer, T. Wang, M. Michaelos, M. Pachitariu, Cellpose: a generalist algorithm for cellular segmentation. Nat. Methods. 18, 100–106 (2021).

98. TN_Ancillary_RNAscope Multiplex Fluorescent V2_ISH-IF Co-Detection Workflow (ICW), (available at https://acdbio.com/search/site/%252Amk%252051-150%252A/cms/support).

99. RNAscope® Multiplex Fluorescent v2 Assay for Cultured Adherent Cells in 96-well Plate Format, (available at https://acdbio.com/system/files_force/SOP45-009A_TechNoteMuxFL_CulturedCells_96well-plateV2_10112017.pdf?download=1).

100. C. A. Koczor, K. M. Saville, J. F. Andrews, J. Clark, Q. Fang, J. Li, R. Q. Al-Rahahleh, M. Ibrahim, S. McClellan, M. V. Makarov, M. E. Migaud, R. W. Sobol, Temporal dynamics of base excision/single-strand break repair protein complex assembly/disassembly are modulated by the PARP/NAD+/SIRT6 axis. Cell Rep. 37, 109917 (2021).

101. D. Hanisch, A. Krumm, T. Diehl, C. M. Stork, M. Dejung, F. Butter, E. Kim, W. Brenner, G. Fritz, T. G. Hofmann, W. P. Roos, Class I HDAC overexpression promotes temozolomide resistance in glioma cells by regulating RAD18 expression. Cell Death Dis. 13 (2022), doi:10.1038/s41419-022-04751-7.

102. M. Guttman, G. N. Betts, H. Barnes, M. Ghassemian, P. Van Der Geer, E. A. Komives, Interactions of the NPXY microdomains of the low density lipoprotein receptor-related protein 1, 5016–5028 (2009).

103. A. L. McCormack, D. M. Schieltz, B. Goode, S. Yang, G. Barnes, D. Drubin, J. R. Yates, Direct Analysis and Identification of Proteins in Mixtures by LC/MS/MS and Database Searching at the Low-Femtomole Level. Anal. Chem. 69, 767–776 (1997).

104. MyAssays Online, (available at https://www.myassays.com/).

