## Supplemental Figures and Materials Table for "AAV Kills Dividing Cells by Depleting PARP1 and Other DNA Damage Response Proteins"

### Materials and Methods:

| Key Resources Table | Source | Identifier | Notes |
|---|---|---|---|
| **Biological Resources** | | | |
| **Cell lines** | | | |
| H9 NPCs | WiCell | hPSCReg ID: WAe009-A | Used to generate hNPCs |
| U2OS | ATCC | Cat# HTB-96 | Used to generate U2OS-LivePAR cells |
| CW468 GSCs | derived from Duke University Patient Samples | Patient samples were collected following IRB-approved protocol 090401 | |
| **Virus** | | | |
| AAV1-CAG-flex-EGFP | Addgene | Cat# 51502-AAV1 | |
| AAV9-CAG-GFP | Addgene | Cat# 37825-AAV9 | |
| AAV9-ΔBC-ITR-CAG-GFP | Salk Viral Vector Core | | |
| AAV9-WT-ITR-CAG-GFP | Salk Viral Vector Core | | |
| **Reagents** | | | |
| Hemocytometer | Fisher Sci | Cat# 0267151B | |
| Tissue Culture Plate 6 wells, Flat Bottom | Fisher Sci | Cat# FB012927 | |
| Nicotinamide ribose | TruNiagen, isolated from 300mg capsules | Tru Niagen® 300mg, no product number listed | |
| 400 mesh copper grids | TedPella | Cat# 01754- F | |
| Uranyl acetate | Ladd research | Cat# 23620 | |
| Whatman 1 filter paper | Whatman | Cat# 1001-085 | |
| C18 desalting columns | Thermo Scientific | Cat# PI-87782 | |
| DNAse I | Thermo Scientific | Cat# EN0521 | |
| PrimeTime Gene Expression Master Mix | Integrated DNA Technologies (IDT) | Cat# 1055772 | |

| | | | |
|---|---|---|---|
| EGFP TaqMan Assay | ThermoFisher Scientific | Assay ID: Mr04329676_mr | cat.# 4331182; 20×, FAM-MGB |
| KpnI-HF | NEB | Cat# R3142S | |
| AAV9 Titration ELISA kit | Progen | Cat# PRAAV9 | Lot#A4004 |
| 96-well Clear Flat Bottom TC-treated Microplate, Sterile | Corning | Cat# 3585 | |
| Falcon® 100 mm TC-treated Cell Culture Dish, Sterile | Corning | Cat# 353003 | |
| µ-Slide 18 Well plates | Ibidi | Cat# 81811 | |
| NuPAGE 4-12% Bis-Tris Gel | Invitrogen | Cat# NP0321BOX | |
| Pre-Cut Nitrocellulose Membranes | Thermo Scientific | Cat# 77010 | |
| Pierce™ BCA Protein Assay Kit | Thermo Scientific | Cat# 23225 | |
| LDS Sample Buffer (4X) | Invitrogen | Cat# NP0007 | |
| SuperSignal West Pico PLUS Chemiluminescent Substrate | Thermo Scientific | Cat# 34580 | |
| PageRuler Plus Prestained Protein Ladder | Thermo Scientific | Cat# 26619 | |
| Dynabeads MyOne Streptavidin T1 | Invitrogen | Cat# 65601 | |
| HCl (hydrochloric acid) | Fisher Sci | Cat# A144-212 | |
| NaOH | Fisher BioReagents | Cat# BP359-500 | |
| Paraformaldehyde (PFA) Granular | Electron Microscopy Sciences | Cat# 19210 | |
| Acetone | Fisher Sci | Cat# A18-1 | |
| Diethyl ether | Fisher Sci | Cat# E134-1 | |
| Ethanol | Fisher Sci | Cat# 2701 | |
| Ethanol | Sigma Aldrich | Cat# E7023 | |
| Methanol | Fisher Sci | Cat# A452SK-4 | |
| Triton X-100 | MilliporeSigma | Cat# 93443 | |
| Triton X-100 | Sigma Aldrich | Cat# X100-5ML | |
| Tween 20 | Sigma Aldrich | Cat# P1379 | |
| NP-40 Surfact-Amps Detergent Solution | Thermo Scientific | Cat# 85124 | |
| Potassium Chloride | Sigma | Cat# P9541 | |
| Sodium Chloride | Sigma Aldrich | Cat# S9888 | |
| Magnesium Chloride | Sigma Life Science | Cat# M4880 | |
| Calcium Chloride Dihydrate | Sigma Life Science | Cat# C7902 | |
| Halt Protease Inhibitor/Phosphatase Inhibitor Cocktail | ThermoFisher | Cat# 78441 | |

| | | |
|---|---|---|
| Protease Inhibitor Cocktail | Sigma-Aldrich | Cat# P8340 |
| Trizma Base | Sigma | Cat# T1503 |
| DL-Dithiothreitol | Sigma Aldrich | Cat# D0632 |
| RIPA Lysis Buffer, 10x | EMD Millipore | Cat# 20-188 |
| Sodium Dodecyl Sulfate | Thermo Scientific | Cat# 28364 |
| Paraformaldehyde, 4% in PBS | Fisher Sci | Cat# J61899.AP |
| PBS (Dulbecco w/o Ca2+ w/o Mg2+, 9.55 g/l) | VWR | Cat# 21-031-CV |
| DPBS/Modified -Ca -Mg | Cytiva | Cat# SH30028.02 |
| DPBS 1x without Ca & Mg | Corning | Cat# 21-031-CV |
| DMEM/F12 | Life Technologies | Cat# 10565 |
| B27 | Life Technologies | Cat# 17504 |
| N2 | Life Technologies | Cat# 17502 |
| Y-27632 | Tocris | Cat# 1254 |
| Corning Costar Ultra-Low Attachment Microplates | Corning | Cat# 3471 |
| Collagenase IV | Gibco | Cat# 17104 |
| mTeSR1 | Stemcell Technologies | Cat# 85850 |
| Matrigel | Corning | Cat# 354230 |
| Noggin | R&D Systems | Cat# 6057-NG |
| SB431542 | Tocris | Cat# 1614 |
| Poly-L-ornithine | Sigma | Cat# P3655 |
| Laminin Mouse Protein | Natural Life Technologies | Cat# 23017015 |
| Heat-inactivated Fetal bovine serum | Bio-Techne | Cat# S11150H |
| Penicillin/streptomycin | Thermo Fisher Scientific | Cat# 15140–122 |
| DMEM | Corning | Cat# 15–017-CV |
| L-glutamine | Thermo Fisher Scientific | Cat# 25030–081 |
| Neurobasal | Gibco | Cat# 12349015 |
| B27 | Gibco | Cat# 12-587-010 |
| Sodium pyruvate | Gibco | Cat# 11360-070 |
| Glutamine | Gibco | Cat# 35050079 |

| | | |
|---|---|---|
| EGF | StemCell Technologies | Cat# 78006 |
| Pen/strep | Gibco | Cat# 15140122 |
| UltraPure Distilled Water | Invitrogen | Cat# 10977-015 |
| DMEM/F12 + GlutaMAX | Gibco | Cat# 10565-018 |
| N-2 Supplement | Gibco | Cat# 17-502-048 |
| B-27 Supplement (50X), minus vitamin A | Gibco | Cat# 12-587-010 |
| Recombinant Human FGF basic/FGF2/bFGF aa TC Grade, CF | Bio-Techne | Cat# 4144-TC-01M |
| Cultrex Reduced Growth Factor Basement Membrane Extract, PathClear | Bio-Techne | Cat# 3433-005-01 |
| Accutase | Innovative Cell Technologies | Cat# AT 104 |
| Propidium Iodide | Sigma-Aldrich | Cat# P4170 |
| 1M TRIS-HCL pH 7.5 | Teknova | Cat# T1075 |
| EDTA | EMD Millipore Corp | Cat# 324503-100gm |
| EDTA | ThermoFisher | Cat# J60877.K3 |
| RapiGest SF reagent | Waters Corp | Cat# 186002122 |
| TCEP (Tris (2-carboxyethyl) phosphine | ThermoFisher | Cat# PI20491 |
| Iodoacetamide | ThermoFisher | Cat# AC122270050 |
| Trypsin | ThermoFisher | Cat# PI90059PM |
| HCl | Fisher | Cat# 60-007-48 |
| BCA Assay | ThermoFisher | Cat# PI23225 |
| 1.7-µm C18 (130) BEHTM beads | Waters Corporation | Cat# 186004971 |
| Acetonitrile | Fisher | Cat# A955-500 |
| Formic Acid | Millipore | Cat# 186004971 |
| Bovine Serum Albumin (BSA) | RPI research Products International | Cat# A30075-100.0 |
| Bovine Serum Albumin (BSA) | Roche | Cat# 3117332001 |
| Horse Serum | Millipore Sigma | Cat# H1270 |
| Dry Milk Powder | Research Products International | Cat# M17200-500.0 |
| Glycerol | Reagents | Cat# C1210500-4C |

| | | | |
|---|---|---|---|
| VECTASHIELD Antifade Mounting Medium with DAPI, 10mL | Fisher Sci | Cat# NC1695563 | |
| Immu-Mount | Epredia | Cat# 9990402 | |
| DMSO (dimethyl sulfoxide) | Fisher Sci | Cat# 3176 | |
| DMSO (dimethyl sulfoxide) | Sigma-Aldrich | Cat# D2650 | |
| Bacteriostatic 0.9% Sodium Chloride | Pfizer | Cat# 00409196607 | |
| MNNG | Molport | Cat# N493990 | |
| NRH | NuChem Sciences | | |
| PDD00017273 | MilliporeSigma | Cat# SML1781 | |
| ABT-888 | Tocris | Cat# 7026 | |
| ABT-888 | Selleck hem | Cat# S1004 | |
| BML-277 | APExBIO | Cat# B1236 | |
| Clear nail polish | O.P.I. nature strong | | |
| RNA-Protein Co-Detection Ancillary kit | ACD Bio | Cat# 323180 | |
| RNAscope H2O2 and Protease Reagents | ACD Bio | Cat# 322381 | |
| RNAscope Multiplex Fluorescent Reagents Kit v2 | ACD Bio | Cat# 323100 | |
| RNAscope Multiplex TSA Buffer | ACD Bio | Cat# 322809 | |
| RNAscope 50X Wash Buffer | ACD Bio | Cat# 310091 | |
| RNAscope Probe EGFP-sense | ACD Bio | Cat# 409971 | |
| Opal 520 Reagent | Akoya Biosciences | Cat# OP-001001 | 1:750 dilution used for RNAscope |
| **Antibodies** | | | |
| DAPI | Sigma Aldrich | Cat# D9542 | 1:10000 dilution used for ICC |
| Phospho-Chk1 (Ser345) (133D3) Rabbit mAb | Cell Signaling Technologies | Cat# 2348 | 1:100 dilution used for ICC, 1:1000 dilution used for Western Blotting |
| Phospho-Chk2 (Thr68) Antibody | Cell Signaling Technologies | Cat# 2661 | 1:100 dilution used for ICC |
| Phospho-Chk2 (Thr68) (C13C1) Rabbit mAb | Cell Signaling Technologies | Cat# 2197 | 1:1000 dilution used for Western Blotting |

| | | | |
|---|---|---|---|
| Anti-ATM (phospho S1981) antibody [EP1890Y] | Abcam | Cat# ab81292 | 1:250 dilution used for ICC, 1:2000 dilution used for Western Blotting |
| ATR (phospho Thr1989) antibody | GeneTex | Cat# GTX128145 | 1:500 dilution used for ICC, 1:1000 dilution used for Western Blotting |
| Phospho-Histone H2A.X (Ser139) (20E3) Rabbit mAb | Cell Signaling Technologies | Cat# 9718 | 1:400 dilution used for ICC, 1:1000 dilution used for Western Blotting |
| 53BP1 Antibody - BSA Free | Novus Biologicals | Cat# NB100-304 | 1:1000 dilution used for ICC, 1:500 dilution used for western blotting, 1:500 dilution used for RNAscope |
| Phospho-PNK1 (Ser114/Thr118) Antibody | Cell Signaling Technologies | Cat# 3522 | 1:50 dilution used for ICC, 1:1000 dilution used for Western blotting |
| Anti-Topoisomerase I-DNA Covalent Complexes Antibody, clone 1.1A | Millipore | Cat# MABE1084 | 1:200 dilution used for Western Blotting |
| Mouse monoclonal anti-LIG3 [1F3] | GeneTex | Cat# GTX70143 | 1:500 dilution used for Western Blotting |
| PARP1 | Proteintech | Cat# 13371-1-AP | 1:1000 dilution used for Western Blotting |
| phospho-XRCC1 [p Ser485, p Thr488] | Novus Biologicals | Cat # NB100-541 | 1:1000 dilution used for Western Blotting |
| GAPDH (14C10) | Cell Signaling Technology | Cat# 2118 | 1:5000 dilution used for Western Blotting |
| Donkey anti-Rabbit IgG (H+L) Highly Cross-Adsorbed Secondary Antibody, Alexa Fluor™ 568 | Invitrogen | Cat# A10042 | 1:250 dilution used for ICC, 1:125 dilution used for RNAscope |

| | | | |
|---|---|---|---|
| Peroxidase AffiniPure™ Donkey Anti-Rabbit IgG (H+L) | Jackson Immunoresearch | Cat# 711-035-152 | 1:5000 dilution used for Western Blotting |
| Peroxidase AffiniPure™ Donkey Anti-Mouse IgG (H+L) | Jackson Immunoresearch | Cat# 715-035-150 | 1:5000 dilution used for Western Blotting |
| **DNA Sequences** | | | |
| Wild type AAV2: AGGAACCCCTAGTGATGGAGTTGGCCACTCCCTCT CTGCGCGCTCGCTCGCTCACTGAGGCCGGGCGACC AAAGGTCGCCCGACGCCCGGGCTTTGCCCGGGCG GCCTCAGTGAGCGAGCGAGCGCGCAGAGAGGGAG TGGCCAA | IDT 20 nmol Ultramer™ DNA Oligo, Biotinylated at 5' end | | |
| Scrambled wild type AAV2: AACAACCTGACGGTACCGCGCGCGTCGCCGTTTCC TGGCGCTGTCGGAGGCGGCAGCACGGTCCAGGAC GGAATCGCTTTGGCGCCCCTAGCGGCAGCGGGGGC GGGACTGATCCAACCGGTACTGCGGTGCACGACGC ACGACC | IDT 20 nmol Ultramer™ DNA Oligo , Biotinylated at 5' end | | |
| Random 145 bp: CCACATACCGTCTAACGTACGGATTCCGATGCCCAG ATATATAGTAGATGTCTTATTTGTGGCGGAATAGCGC CAGAGCGTGTAGGCCAACCTTAGTTCTCCATGGAA GGCATCTACCGAACTCGGTTGCGCGGCCAAATTGG AT | IDT 20 nmol Ultramer™ DNA Oligo , Biotinylated at 5' end | | |
| ΔBC: CCACATACCGTCTAACGTACGGATTCCGATGCCCAG GAACCCCTAGTGATGGAGTTGGCCACTCCCTCTCT GCGCGCTCGCTCGCTCACTGAGGCCGCCCCGCGGC CTCAGTGAGCGAGCGAGCGCGCAGAGAGGGAGTG GCCAA | IDT 20 nmol Ultramer™ DNA Oligo , Biotinylated at 5' end | | |
| **Software and Algorithms** | | | |
| Image J | Image J version 1.54g ImageJ version 1.54f | https://wsr.imagej.net/distros/osx/ij154-osx-java8.zip | |

| | | |
|---|---|---|
| Cell Pose | Carsen Stringer and Marius Pachitariu | version 3.1.1.1 |
| Cell Profiler | Cimini Lab at the Broad Institute of MIT and Harvard | version 4.2.8 |
| FlowJo | BD Biosciences | version 11 |
| Prism | GraphPad Software | version 10.4.2 |
| PeaksStudio 8.5 | Bioinformatics solutions Inc. | |
| **Cell Culture Media and Solutions** | **Ingredients** | |
| N2B27 Media | DEMEM F12 (Life Technologies/Gibco), 1X N2 (Life Technologies/Gibco), 1X B27 (Life Technologies Supplements/Gibco) | |
| U2OS Media | DMEM (Corning), 1X L-glutamine (Thermo Fisher Scientific), 10% heat inactivated fetal bovine serum (Bio-Techne), and penicillin/streptomycin (Thermo Fisher Scientific) | |
| GSC media | Neurobasal media (Gibco), 1X B27 (Gibco), 1X sodium pyruvate (Gibco), 1X glutamine (Gibco), EGF (40ng/mL; StemCell Technologies), FGF (40ng/mL; Bio-techne), and penicillin/streptomycin (Gibco) | |
| TBS | 250mM trizma base (Sigma), 1.5M NaCl (Sigma Aldrich), 25mM KCl (Sigma), HCl to pH 7.4 (Teknova) in water | |
| Blocking buffer for immunocytochemistry | 3% horse serum (Millipore Sigma), 0.25% Triton X-100 (Millipore Sigma) in TBS | |
| 1X RIPA Buffer | 100mM DTT (Sigma Aldrich), 10% SDS (Thermo Scientific), 1X Protease inhibitor (Sigma-Aldrich) in 1X RIPA (EMD Millipore) | |
| Swelling buffer for pull down | 10 mM Tris-HCl (Teknova), 2 mM MgCl2 (Sigma Life Science), 3 mM CaCl2 (Sigma Life Science) in ultrapure water (Invitrogen). | |
| Lysis buffer for pull down | 10% glycerol (Reagents) in swelling buffer | |
| NP40 lysis buffer for pull down | 150 mM NaCl; Sigma Aldrich, 1% NP-40; Thermo Scientific, 50 mM Tris-HCl | |
| TNE buffer for mass spectrometry | 50 mM Tris pH 8.0, 100 mM NaCl, 1 mM EDTA (ThermoFisher) | |
| Buffer A for mass spectrometry | 98% H2O, 2% ACN (Fisher), 0.1% formic acid (Millipore) | |

| Buffer B for mass spectrometry | 100% ACN (Fisher), 0.1% formic acid (Millipore) |
|---|---|

### Supplementary Figures:

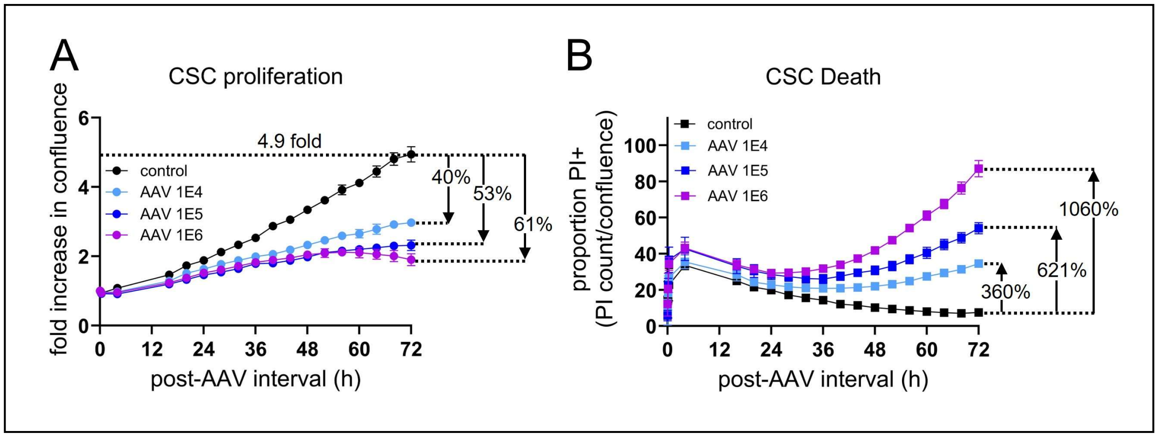

**Figure S1. Recombinant AAV induces dose-dependent toxicity in patient-derived glioblastoma stem cells (GSCs)** (**A**) Example cell confluence measurements from time-lapse microscopy of GSCs plated on a 96-well plate and infected with rAAV at various MOIs and saline control demonstrate dose-dependent attenuation of cell proliferation. Each point represents normalized confluence averaged over n = 3 wells. (**B**) Dose-dependent rAAV-induced cell death is measured in the same wells by calculating the proportion of PI+ GSCs (n = 3 wells).

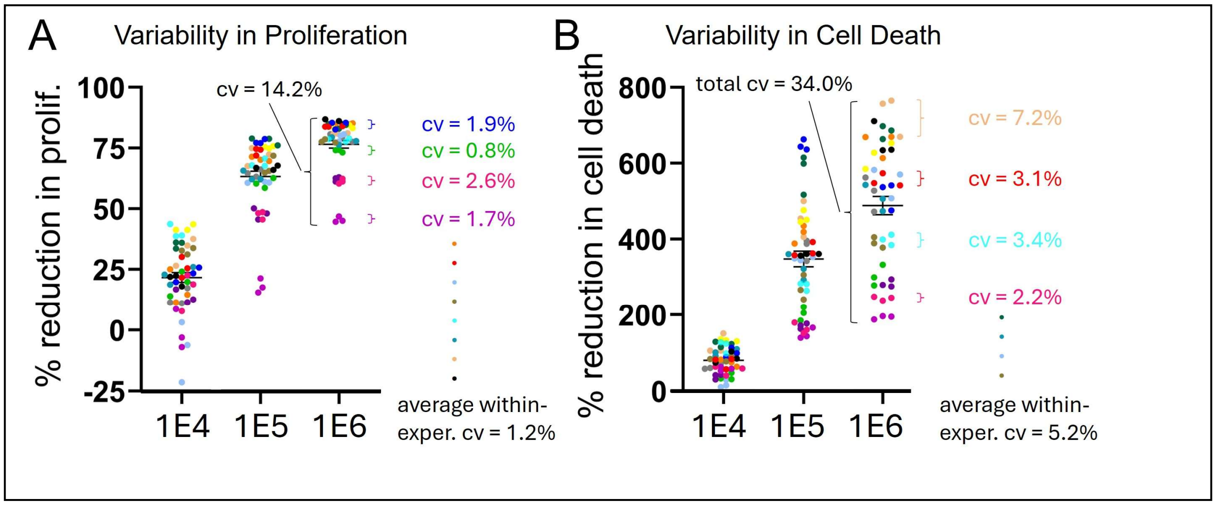

**Figure S2. Variability in rAAV toxicity is substantially smaller within experiments than between experiments** (**A**) Data from Figure 1D for reduction in cell proliferation and (**B**) Figure 1E increase in cell death at 72 hours are replotted such that all wells within a given experiment are marked with the same color (n = 16 experiments) to compare the variability among wells within the same experiments to the variability between different experiments. This comparison is quantified by calculating the total coefficient of variation (cv) over all 48 wells at a given MOI and comparing it to the within-experiment cv averaged over all 16 experiments. See Results for additional details.

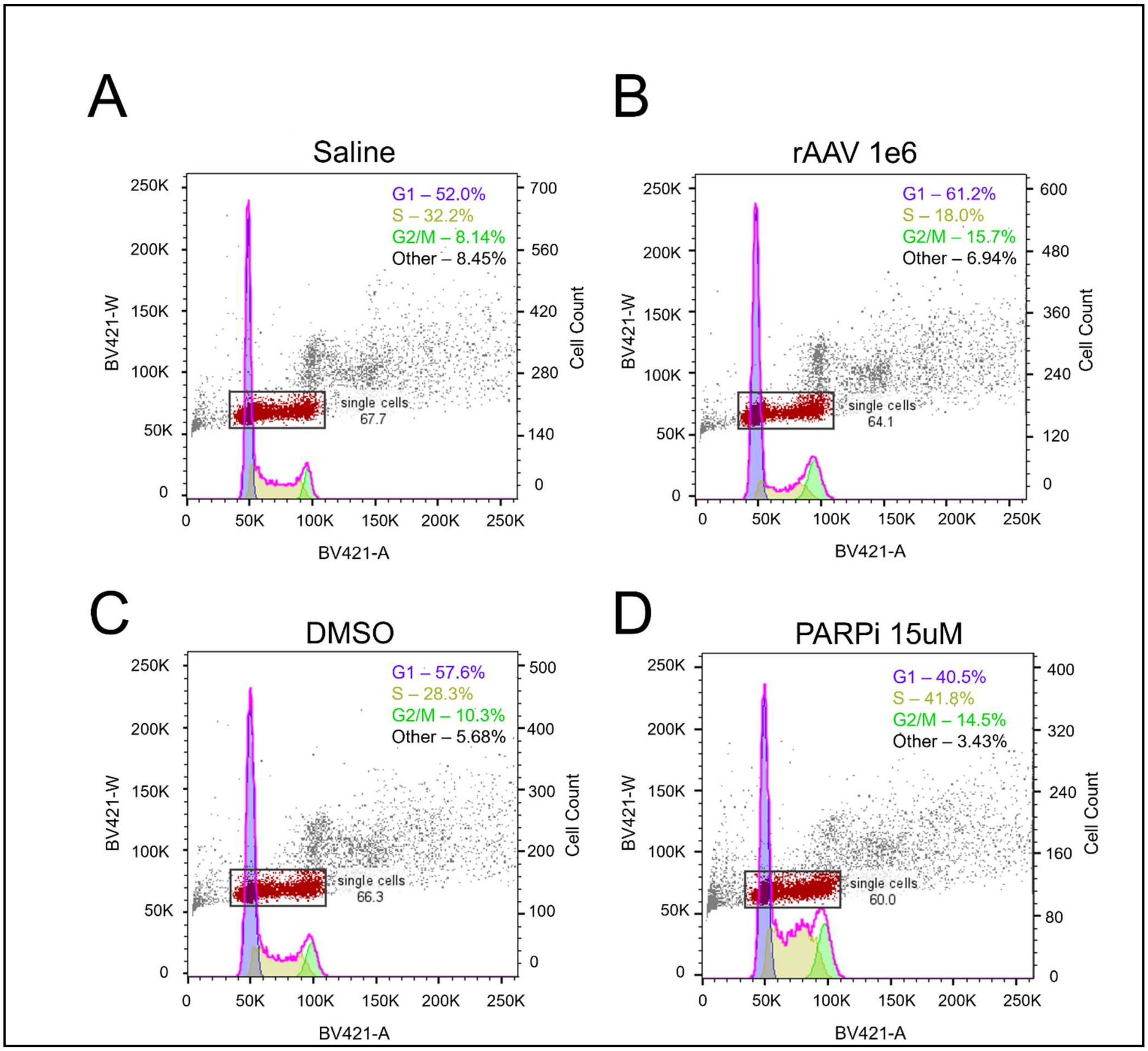

**Figure S3. HNPCs treated with rAAV or PARP inhibitor do not exhibit substantial cell cycle arrest.** FACS cell cycle analysis of hNPCs 24 hours after treatment with (**A**) saline, (**B**) rAAV (MOI = 1E6 gc/ml), (**C**) DMSO, or (**D**) PARP inhibitor with DMSO vehicle (ABT-888, 15uM) are stained for DAPI is quantified for DNA content. Scatter plots show the width (BV421-W) versus area (BV421-A) for acquired events. Events determined to be single cells are gated and shown in red with single cell percentages displayed. The same gate area and position are used for all samples in (**A**)-(**D**)). Overlayed histograms and plotted with the fraction of cells in each phase determined using FlowJo Cell Cycle function with Watson (Pragmatic) model and default settings. G1 is shown in purple, S is shown in yellow, G2/M is shown in green, and the summation of all phases is shown in pink. The fraction of single cells within each cell cycle is annotated on the histogram and does not exhibit substantial cell cycle arrest.

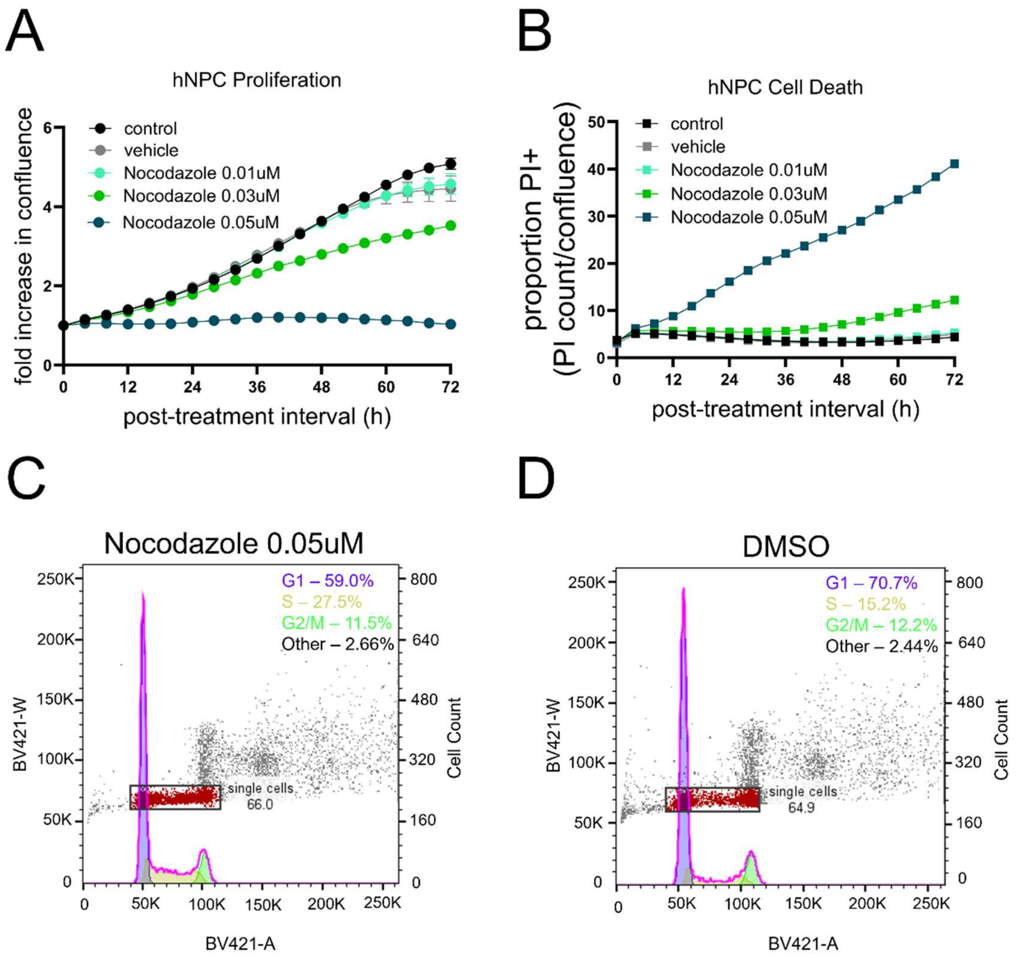

**Figure S4. Treatment with Nocodazole induces dose-dependent toxicity but does not induce substantial cell cycle arrest in hNPCs.** (**A**) Example of time lapse microscopy measurement of hNPCs treated with Nocodazole at various doses compared to saline and vehicle control demonstrate a strong dose-dependent attenuation of cell proliferation and (**B**) cell death (n = 3 wells per condition). FACS cell cycle analysis of hNPCs 24 hours after treatment with (**C**) DMSO, or (**D**) Nocodazole (0.05uM) stained for DAPI and quantified for DNA content shows no clear evidence of cell cycle arrest. Cell cycle plots and analysis were generated as described in Figure S3.

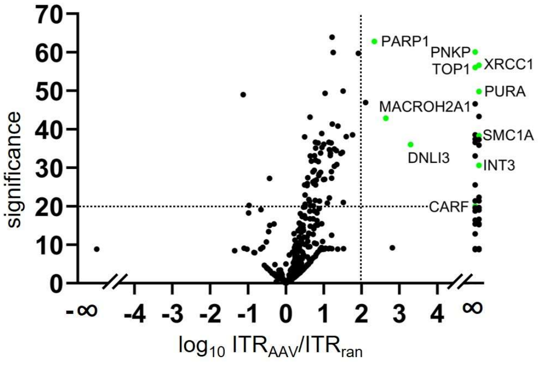

**Figure S5. Affinity pull-down assay of AAV2 ITR in mouse NPCs** (**A**) Relative abundance of nuclear proteins isolated by AAV2 ITR ("ITRAAV") compared to a random 145-base pair single-stranded DNA sequence ("ITRRAN") via an affinity pull-down assay in primary mouse NPCs and measured by mass spec. Nineteen of the top 23 proteins (defined as significance ≥ 20, log$_{10}$(ITRAAV/ITRRAN) ratio ≥ 2) significance > 20, log$_{10}$(ITRAAV/ITRRAN) ratio ≥2) pulled down by ITRAAV were not detectable in the ITRRAN fraction, yielding a log$_{10}$(ITRAAV/ITRRAN) ratio = ∞. Ten of these top proteins were identified by mass spec in both glioblastoma stem cells (Fig. 3) and mouse NPCs and are marked by green points and labeled.

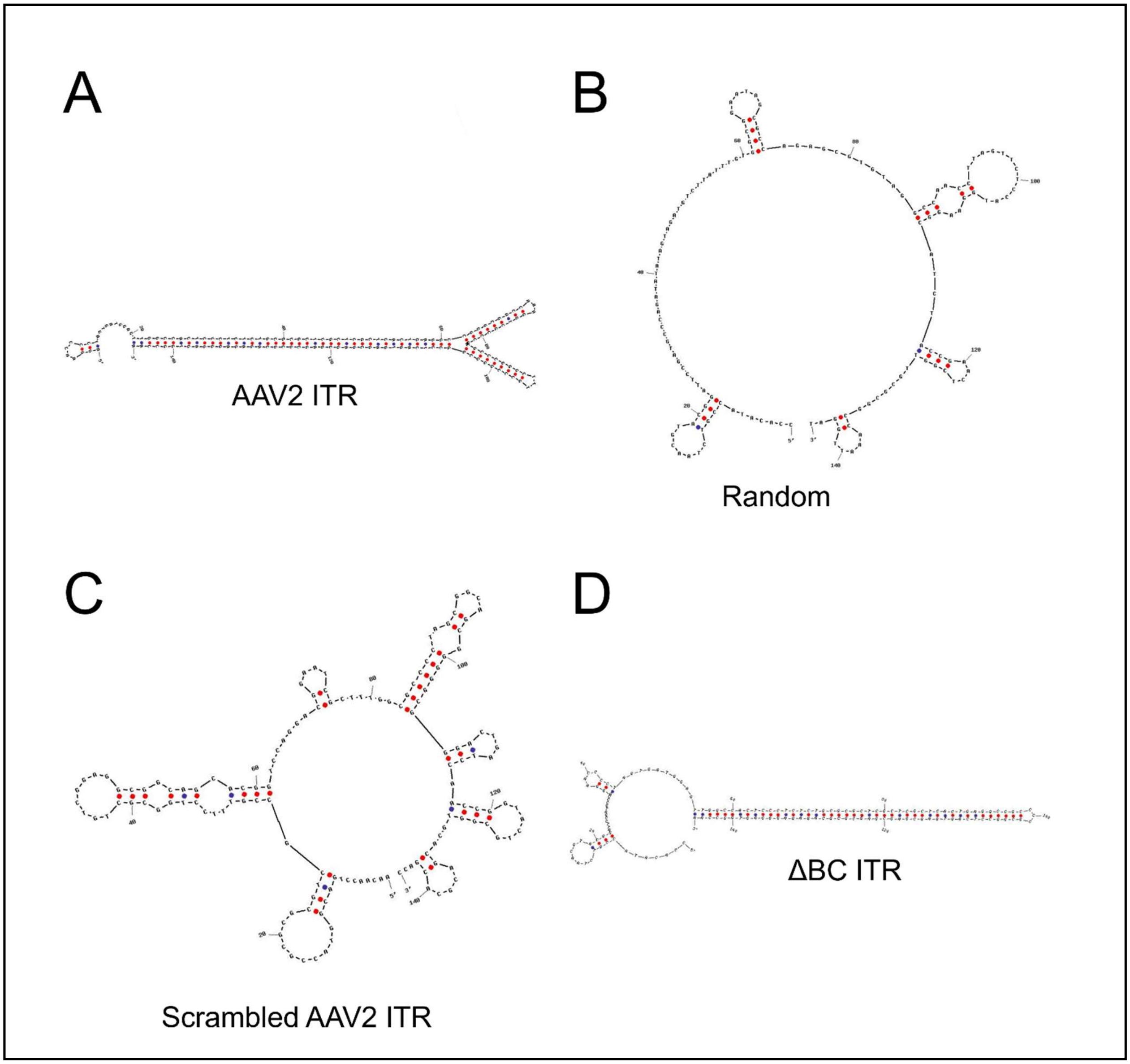

**Figure S6. Secondary structure model of** (**A**) AAV2 ITR (145bp), ΔG=-82.837 kcal/mol (**B**) Random 145bp oligomer, ΔG=-15.729 kcal/mol (**C**) Scrambled AAV2 ITR oligomer (145bp), ΔG=-25.498 kcal/mol and (**D**) ΔBC ITR (145bp), , ΔG=-70.033 kcal/mol. The random 145bp oligomer is contains equal numbers of each of the 4 nucleotides. The ΔBC ITR contains the original 111bp ΔBC ITR sequence[81] with an additional 34 base pairs appended to the 5' end, which were taken from the 5' end of the random 145bp oligomer to control for effects of DNA length. Structures were generated using the mFold algorithm (through IDT's OligoAnalyzer hairpin settings with SpecSheet Parameter Sets: Oligo Conc 0.25uM, $Na^+$ Conc 50mM, $Mg^{++}$ Conf 0mM, dNTPs Conc 0mM, and DNA Target Type). For each sequence, the most stable output was selected.

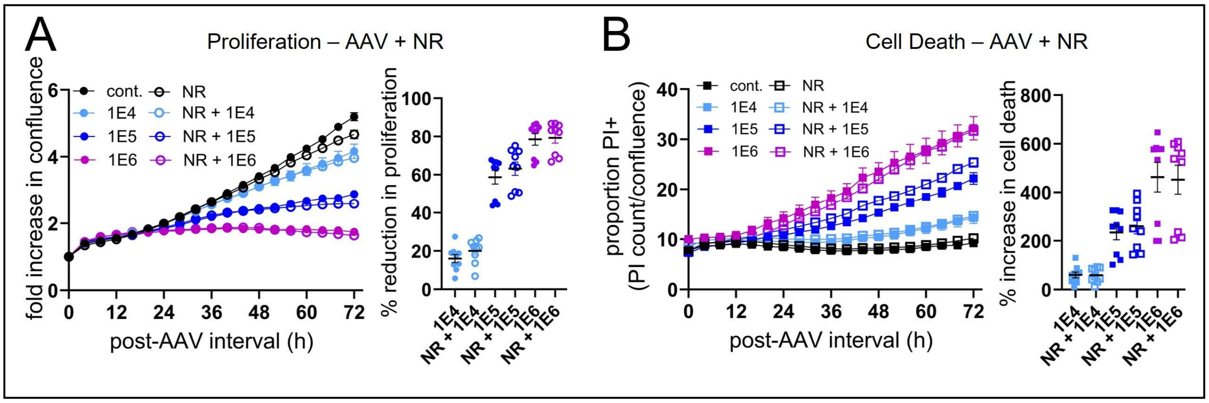

**Figure S7. rAAV toxicity is not rescued by NR supplementation.** (**A**) Example cell confluence measurements from time-lapse microscopy of hNPCs infected with rAAV at various MOIs with (closed points) and without (open points) NR supplementation demonstrate no rescue of rAAV-induced attenuation of proliferation or (**B**) rAAV-induced cell death.

| Accession | Name(s) | DDR Citation | Interacts w/ PARP1? | PARP1 Citation | Log Ratio | Significance |
|---|---|---|---|---|---|---|
| **P09874** | PARP1 | PMC4936651 | N/A | | ∞ | 84.3 |
| **P49916** | LIG3 (DNLI3) | PMC4333375 | Y | PMC9723622 | ∞ | 52.7 |
| **P18887** | XRCC1 | PMC4333375 | Y | PMC9723622 | ∞ | 48.4 |
| **Q96T60** | PNKP | PMC4333375 | Y | PMC5389470 | ∞ | 47.1 |
| **P12956** | XRCC6 (KU70) | PMC3589933 | Y | PMC2861645 | ∞ | 46.6 |
| **Q9Y265** | RUVB1 (RUVBL1) | PMC11015013 | Y | PMC4177365 | ∞ | 40.1 |
| **O75367** | H2AY (MACROH2A1) | PMC6908255 | Y | PMC6908255 | ∞ | 39.3 |
| P40429 | RL13A (RPL13A) | | | | ∞ | 38.7 |
| **P59998** | ARPC4 | PMC6145447 | | | ∞ | 37.5 |
| **P20700** | LMNB1 | PMC8397269 | | | ∞ | 37.0 |
| **P78347** | GTF2I (TFII-I) | PMC3859932 | Y | PMC2493457 | ∞ | 36.6 |
| **O00159** | MYO1C (NM1) | PMC7066169 | | | ∞ | 36.6 |
| **Q9NXV6** | CARF (CDKN2A) Interacting Protein | PMC4140265 | | | ∞ | 36.3 |
| **P11387** | TOP1 | PMC8095074 | Y | PMC8095074 | ∞ | 36.2 |
| **Q00577** | PURA | PMC2586959 | | | ∞ | 24.2 |
| Q8N1F7 | NUP93 | | | | ∞ | 22.9 |
| **Q9NTI5** | PDS5B | PMC8198109 | | | ∞ | 22.8 |
| Q13045 | FLII | | | | ∞ | 22.6 |
| **Q9NZC9** | SMAL1 (SMARCAL1) | PMC2791023 | Y | PMC9794062 | ∞ | 22.4 |
| P67936 | TPM4 | | | | ∞ | 22.2 |
| **Q9H0A0** | NAT10 | PMC10591715 | Y | PMC9389688 | ∞ | 22.1 |
| Q92747 | ARC1A (ARPC1A, SOP2L) | | | | ∞ | 22.0 |
| **O43795** | MYO1B | PMC6051730 | | | ∞ | 21.9 |
| P55795 | HNRH2 (HNRNPH2) | | | | ∞ | 21.7 |
| **Q68E01** | INT3 (INTS3) | PMC2762097 | | | ∞ | 21.7 |
| Q9UEG4 | ZN629 (ZNF629) | | | | ∞ | 21.6 |
| **P14618** | KPYM (PKM) | PMC5752508 | | | ∞ | 21.5 |
| **Q16531** | DDB1 | PMC2894263 | Y | PMC6591728 | ∞ | 21.4 |
| **P0DMV9** | HS71B (HSP70-1B, HSPA1B) | PMC2728274 | Y | PMC2728274 | ∞ | 21.1 |

| | | | | | | |
|---|---|---|---|---|---|---|
| P30876 | RPB2 (POLR2B) | | | | ∞ | 21.0 |
| **Q9P2D1** | CHD7 (Chromodomain Helicase DNA Binding Protein 7) | PMC7666215 | Y | PMC7666215 | ∞ | 20.6 |
| **P52292** | IMA1 (KPNA2, SRP1-alpha) | PMC6600173 | | | ∞ | 20.1 |

**Table S1. Mass spectrometry results from an affinity pull-down experiment of nuclear proteins isolated from patient-derived glioblastoma stem cells with AAV2 ITR compared to a random 145-base pair single-stranded DNA sequence.** Results represent the top 32 proteins (with significance ≥ 20, $\log_{10}(ITR_{AAV}/ITR_{RAN})$ ratio ≥ 2) pulled down by AAV2 ITR. Twenty-four of these proteins participate in DDR are indicated in red. PMCID numbers for citations describing their role in DDR and interaction with PARP1 are listed.

| Accession | Name(s) | Log Ratio | Significance |
|---|---|---|---|
| **Q60596** | XRCC1 | ∞ | 60.0 |
| **Q9JLV6** | PNKP | ∞ | 56.6 |
| **Q04750** | TOP1 | ∞ | 56.0 |
| **P42669** | PURA | ∞ | 49.8 |
| O35295 | PURB | ∞ | 46.6 |
| Q99JF8 | PSIP1 | ∞ | 43.3 |
| P60867 | RS20 | ∞ | 38.5 |
| **Q9CU62** | SMC1A | ∞ | 38.3 |
| P51612 | XPC | ∞ | 37.4 |
| Q6A026 | PDS5A | ∞ | 37.3 |
| Q8BJ37 | TYDP1 | ∞ | 36.6 |
| Q9Z0H3 | SNF5 | ∞ | 35.8 |
| Q9DBG6 | RPN2 | ∞ | 33.1 |
| **Q7TPD0** | INT3 (INTS3) | ∞ | 30.6 |
| Q9CW03 | SMC3 | ∞ | 25.5 |
| B2RY56 | RBM25 | ∞ | 22.3 |
| Q7TQC5 | APTX | ∞ | 21.4 |
| P33174 | KIF4 | ∞ | 21.4 |
| **Q8BI72** | CARF | ∞ | 20.0 |
| **P97386** | LIG3 (DNLI3) | 3.292923761 | 36.0 |
| **Q9QZQ8** | H2AY (MACROH2A1) | 2.638018914 | 42.8 |
| **P11103** | PARP1 | 2.337321848 | 62.8 |
| Q61990 | PCBP2 | 2.103070322 | 46.9 |

**Table S2. Mass spectrometry results from an affinity pull down experiment of nuclear proteins isolated from mouse NPCs (E15.5 mice) with AAV2 ITR compared to a random 145-base pair single stranded DNA sequence.** Results represent the top 23 proteins (with significance ≥ 20, $\log_{10}(ITR_{AAV}/ITR_{RAN})$ ratio ≥ 2) pulled down by AAV2 ITR. Nineteen of these 23 proteins were undetectable in the random 145-base pair mass spec results. The 10 proteins that were also present in the top candidate AAV2 ITR-binding proteins identified in patient derived glioblastoma stem cells are highlighted in green.
